# Using passive acoustic monitoring to estimate the abundance of a critically endangered parrot, the great green macaw (*Ara ambiguus*)

**DOI:** 10.1101/2022.12.29.519860

**Authors:** Thomas C. Lewis, Ignacio Gutiérrez Vargas, Sam Williams, Andrew P. Beckerman, Dylan Z. Childs

## Abstract

Most conservation relies on being able to estimate population size accurately. The development, implementation and adaptation of effective conservation strategies rely on quantifying the impacts of different threats on population dynamics, identifying species that need conservation management, and providing feedback on the effectiveness of any management actions. However, current approaches are not suitable for wide-ranging species that reside in tropical ecosystems. Here we use the great green macaw *Ara ambiguus* as a case study to show that passive acoustic monitoring is an effective tool for collecting data that can then estimate abundance. We estimate a population of 485.65 ± 61 SE great green macaws in Costa Rica during the breeding, suggesting the population here is larger than previously estimated. We have also highlighted potentially important areas for the species in regions that had not previously been studied. We have demonstrated at a population scale that passive acoustic monitoring (PAM) offers conservationists an efficient and effective way to understand population dynamics. With a high proportion of parrot species threatened globally, passive acoustic monitoring will enable effective monitoring and become an essential tool in conservation planning and evaluation. PAM technology has enormous potential to facilitate such assessments because it is easily scalable, recordings can be stored and re-analysed as machine learning, and abundance estimation techniques become more advanced.

## 1 Introduction

Given the scale of the current threats to wild populations and limited funding, conservation actions need to be efficient and effective. Most conservation relies on being able to estimate population size accurately. This is vital for quantifying population dynamics, the impacts of threats and identifying species that require conserving. Measuring population abundance accurately and efficiently should be a priority. For many animal species, methodologies such as distance sampling, capture-mark-recapture, and simple counts are robust, providing good data to base on- and feedback to conservation management plans. However, highly mobile, group-living species violate key assumptions of these methodologies. For example, one of the critical assumptions of distance sampling, a widely used methodology to estimate population abundance, is the independence of sites and observations. Highly mobile, group-living species can violate assumptions by grouping together and moving between locations (Dénes et al., 2018). Unaccounted for, these factors can cause large estimation errors and mask important ecological patterns (Wenger & Freeman, 2008). Consequently, population estimates required to assess the red list category of highly mobile species, such as the great green macaw (Ara ambiguus), can be statistically weak.

As with other high-throughput sensing technologies, such as remote sensing, LIDAR and camera traps, Passive Acoustic Monitoring (PAM) offers a way to increase the scale and accuracy of wildlife monitoring, overcome the challenges of assessing highly mobile species and reduce the cost and on-the-ground expertise (Gibb et al., 2019). This rapidly expanding field can benefit from developments and cost-reductions in Automated Recording Units (ARUs) (Snaddon et al., 2013; Hill et al., 2019; Teixeira, Maron and Rensburg, 2019) and dramatically increase the quantity of data that can be collected at any given time (Gibb et al., 2019). ARUs provide an efficient and non-invasive way to measure a wide variety of metrics, including community composition (Pillay et al., 2019; Bradfer-Lawrence et al., 2020), individual breeding biology (Marin-Cudraz et al., 2019), occupancy (Wood et al., 2019) and abundance (Marques et al., 2013; Pérez-Granados et al., 2019).

Occupancy-based methods to estimate abundance offer an opportunity to utilise passive acoustic monitoring (PAM) to enable population monitoring to be carried out at scale. One occupancy model that presents the opportunity to estimate abundance is the Royle-Nichols (RN) model for estimating abundance from repeated detection/non-detection surveys (Royle & Nichols, 2003). There are only two studies utilising the RN model with PAM data in the literature, both involving bats (Milchram and Bruckner 2018; Mena et al., 2021). Comparing generalised random encounter models (gREM) and RN models gave similarly reliable estimates of bat population density (Milchram and Bruckner 2018). RN models have also been used for camera trapping studies (e.g. Duquette et al., 2014; Van der Weyde et al., 2018; Rogan et al., 2019) and have been shown to compare well to other techniques, such as radio telemetry, to estimate population abundance (Duquette et al., 2014). When compared to N-mixture time-to-detection models, RN models were also found to have similar performance (Strebel et al., 2021)

The RN model takes advantage of the fact that variation in abundance creates variation in detection probability. Thus heterogeneity in detection probability can be used to estimate the underlying distribution of abundances. It follows, then, that one of the central assumptions of the RN model is that the only source of heterogeneity of detection probability is heterogeneity in abundance. However, in a situation where a population is not closed or individual detection probabilities are not constant. In that case, the unmodeled heterogeneity in capture probability will lead to an overestimation of detection probability and, therefore, an underestimation of abundance (Marques et al., 2013). Heterogeneity in individuals’ detection probabilities could result from intrinsic behavioural variation between individuals within a population or extrinsic factors such as habitat, time, season, weather and temperature (Veech et al., 2016). Extrinsic factors can be addressed by including relevant covariates in the models (Royle & Nichols, 2003). However, intrinsic factors are much more challenging to account for, and for many species, this may present an insurmountable violation of occupancy model assumptions. Another issue with using the RN framework, particularly extreme for camera trapping studies, is that recording devices (camera traps or ARUs) are often put out in locations that increase the possibility of detection. For example, selecting known sites or placing numerous camera traps within known territories means detections are not independent between locations or individuals (Rogan et al., 2019). These issues can be avoided if care is taken to consider the behaviour of the study species in the sampling design.

Parrots are one group of highly mobile, at-risk animals that violate assumptions of traditional monitoring techniques (Casagrande and Beissinger 1997; Dénes et al. 2018) and could be a good fit for combining PAM with RN occupancy modelling. Parrots are one of the most endangered bird families, with 42% being threatened with extinction (IUCN 2020b). Robustly estimating the abundance of parrots is extremely difficult, with most current estimates based on roost counts (e.g. Wright et al., 2018; Zulian et al., 2018), distance sampling (Dénes et al., 2018), expert opinion or a combination (e.g. BirdLife International, 2020). Although parrot behaviour prohibits the use of traditional techniques, their behaviour could be ideally suited to PAM and RN modelling to estimate their abundance. They are loud and social, meaning they frequently call one another, even if they are not nearby.

Two key factors driving parrot and other avian distribution and abundance are nest site and food availability (Loiselle and Blake 1991; Poulin et al. 1992; Cockle et al. 2010). This means local abundance is closely related to the number and quality of available nest sites and the quantity and quality of available food. In addition, many parrots show plasticity in diet and foraging strategies to track seasonal variation in food availability across the landscape (Renton et al., 2015). However, this plasticity in the diet does not match their selection of nest sites. As most parrots are secondary cavity nesters, they rely on naturally forming cavities or for other species to excavate their nest sites (Forshaw 2010). A given parrot species might typically use only one to three tree species as nest sites (Renton et al., 2015), suggesting that breeding season abundance will be tied to their preferred nest tree species (e.g. Oliveira et al., 2021).

During breeding, nest site availability influences parrot distribution (Cockle et al., 2010; de la Parra-Martínez et al., 2015; Renton et al., 2015). Required cavity size is directly related to body size, with larger species requiring larger cavities (Renton et al., 2015). Primary forest has more large cavities than secondary and logged forests (Marsden and Pilgrim 2003; Cockle et al. 2010; de la Parra-Martínez et al. 2015), so we would expect the less disturbed forest to be positively related to breeding season abundance in large parrots. Some large parrots, such as the scarlet macaw (SCM - Ara macao) (Vaughan et al., 2005) and blue-winged macaws (Primolius maracana) (Nunes and Galetti, 2007), can persist in degraded, fragmented landscapes, possibly because these species have a broad nesting niche and generalist diets. Other species, such as the blue-throated macaw (Ara glaucogularis) and hyacinth macaw (Anodorhynchus hyacinthinus) (Oliveira et al. 2021) are habitat specialists and, therefore, are severely impacted by habitat degradation and loss of crucial food and nesting species (Oliveira et al. 2021; Herzog et al. 2021).

The great green macaw (GGM) is one of six critically endangered Neotropical parrots (IUCN 2020b). Their population is estimated to be between 500 and 1000 mature individuals, although it is unclear how robust this estimate is as there are areas that are difficult to assess and no statistically robust methods to estimate abundance therefore, the estimate is based primarily on expert opinion (BirdLife International 2020). The northern limit of their range is the Caribbean slope of southeastern Honduras, where their current population is estimated to be ∼260 mature individuals. There is a subpopulation on the Caribbean slope of southern Nicaragua and north-eastern Costa Rica, with an estimated ∼160 mature individuals (BirdLife International 2020). However, the Macaw Recovery Network carried out a non-breeding season roost count in 2021 and estimated the population to be ∼336 individuals (Macaw Recovery Network 2021). In both Panama and Colombia, they can be found in the Caribbean and Pacific lowlands. There is a lack of data on the Panama population (BirdLife International 2020). However, the population in Parque Nacional Cerro Hoya on the Pacific coast of Panama is thought to number ∼60 individuals (Bolcato pers comms). In Colombia, it is suggested that the population has declined from an estimated 1700 mature individuals to ∼100 mature individuals in a decade. The population in Colombia is fragmented into northwest and southwest, with the southwestern population extending into northwestern Ecuador, which are extremely challenging areas to monitor in. A separate small population of the subspecies guayaquilensis in western Ecuador (Fjeldså et al. 1987) is the southern limit of the species range. The two Ecuadorian populations are not thought to be over 50 mature individuals (BirdLife International 2020).

In Costa Rica, the GGM’s range was reduced by ∼90% by the end of the 1990s (Chassot and Monge 2002) but may have increased slightly since then (Fink et al. 2021). One of the main drivers of this decline was habitat loss and degradation, with the loss of around 90% of the mountain almond (Dipteryx panamensis), a vital food and nesting tree (Monge et al. 2003; Chassot et al. 2007; Monge et al. 2012). In Costa Rica, 85% of the monitored GGM’s nests are in mountain almond (Lewis et al. in prep), and their diet consists of over 80% mountain almonds during the breeding season (Monge et al. 2012). Therefore, our current understanding is that at this time of the year, GGM distribution is closely related to the distribution of the mountain almond, which is mainly found in lowland areas with Ultisol soil type (Chun 2008). At the end of the breeding season (∼May), GGMs disperse and leave the breeding areas. It is currently unclear where they disperse, but it is known that they can be found in large flocks in the foothills of the Cordillera Central from September to October (Bolcato 2020). One previous study has attempted to estimate the number of GGMs in the Costa Rica / Nicaragua population using an extended point count methodology. The study estimated 302.93 ± 513.78 individuals in Costa Rica and 532.45 ± 251.33 in Nicaragua (Monge et al. 2010).

We need robust monitoring methodologies to allow us to develop and sustain effective conservation. Unfortunately, current methodologies are not robust when used to estimate the abundance of highly mobile species in environments such as tropical rainforests. PAM provides a way to address these current challenges, allowing monitoring to be carried out at scale whilst minimising on-the-ground work. To be effective, we must have a modelling approach that can use the data PAM generates. Here we demonstrate that the RN model can estimate the abundance of a challenging study species such as the GGM.

## 2 Methods

### 2.1 Study site and design

The study area is situated in northern Costa Rica in a ∼3000km^2^ region of fragmented Caribbean lowland forest (Fig. 1). This includes a known breeding area (Monge *et al*., 2012, Macaw Recovery Network, *unpub. data*) and areas where the breeding status of GGM is unknown. Land use is split between cattle pasture, pineapple and other annual crops, and primary and secondary forests (Fagan et al., 2013; Jadin et al., 2016; Karra et al., 2021). The annual rainfall is ca. 4667mm (2009-2014), with a drier period between January and April (Gilman et al. 2016).

**Figure 1:**
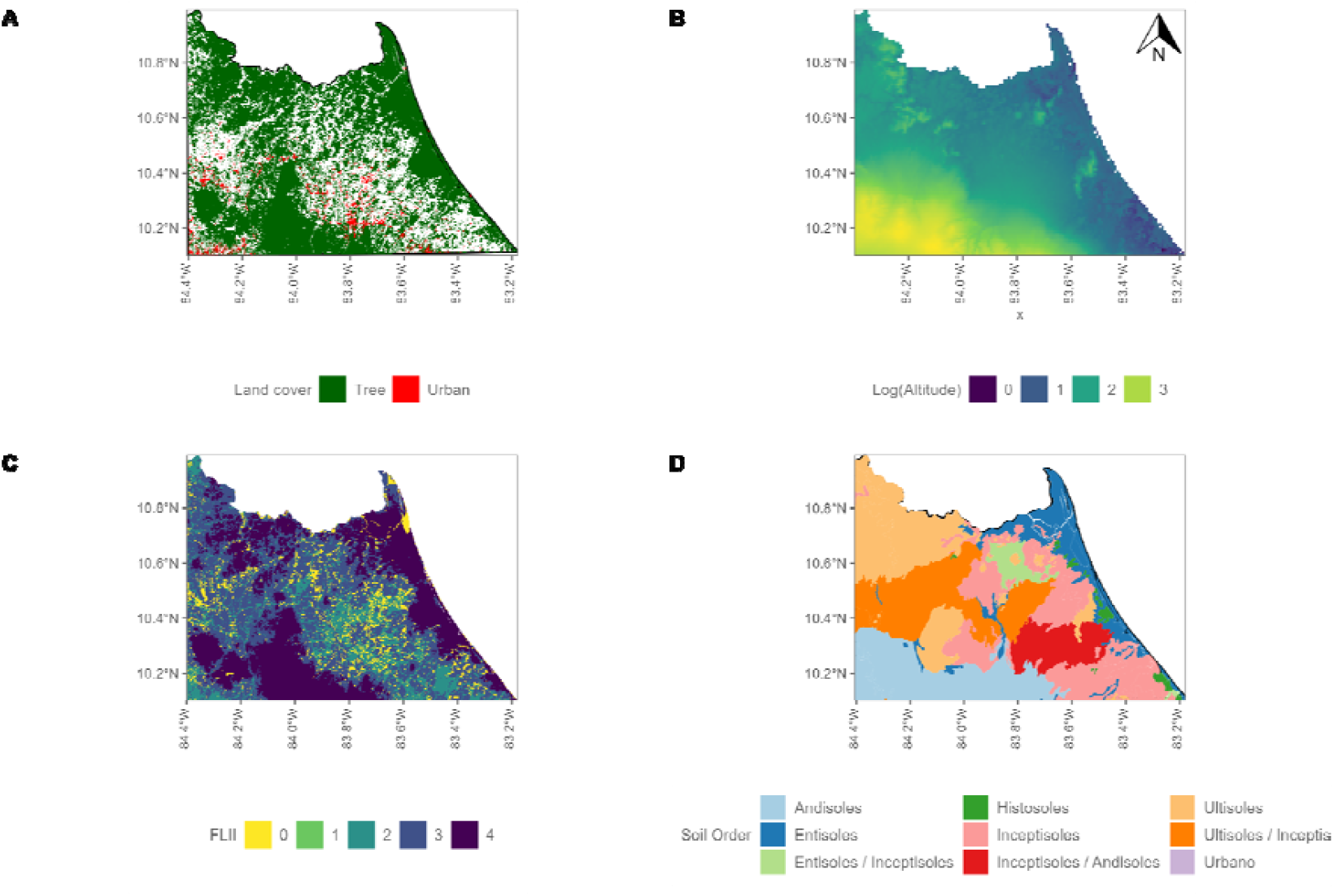
Maps of all the landscape covariates A) Land cover / land use (LCLU) covariates; tree cover and urban cover B) Log altitude C) Forest Landscape integrity index (FLII) D) Soil order.

We conducted the study over two months during the peak 2020 breeding season (Feb-Mar). This period is when breeding pairs forage for food for chicks and before any chicks have fledged the nest (Macaw Recovery Network, *unpub. data*). We placed the ARUs on a 10km grid; we consider this distance large enough to make the locations independent during the study period. This is because a radio-telemetry study found that GGM pairs in Costa Rica maintained breeding ranges of ∼5.5km^2^ (Powell et al., 1999), meaning a 10km grid with each grid square covering 100km^2^ is sufficient to ensure independence. The initial point locations were chosen by taking the centre of every grid square in a 10km grid laid across the study area. Around each point, we used a 1 km buffer where we could move the point if there were accessibility or permission issues with the initial point location. We established forty-three ARU sites following the above principles.

Recordings were captured using AudioMoth 1.1.0 ARUs (Hill *et al*., 2019; LabMaker, Germany). The devices were installed on the tallest accessible tree at each site. There were four 30-minute survey periods (herein “survey periods”) a day (7:00-7:30, 10:00-10:30, 13:00-13:30, 16:00-16:30) between 29 January 2020 and 19 March 2020. Recordings were taken at 32kHz or 48kHz. The sampling frequency was inconsistent as there was an error in configuring some devices for a portion of the deployment period.

### 2.2 Data preparation and statistical analysis

#### 2.2.1 Great green macaw detection data

Detection data were obtained from ARU recordings using a recogniser developed by Lewis *et al*. (2022). This recogniser detects GGM using a two-step pipeline: template matching and supervisor machine learning classification. Multiple templates (n = 6) were used to capture the highly varied call types of the GGMs, and then detections were fed into a random forest (Breiman 2001). The random forest was trained using a spatiotemporally pseudo-random training dataset (Sobol 1967; Antonov and Saleev 1979) that consisted of 8408 GGM and 33129 negative cases. Precision was low (0.56), so all detections of GGMs were manually cleaned to ensure only GGM detections were used for modelling. To derive occupancy data, we pooled and converted detections in each recording into either 1 (detection) or 0 (non-detection).

#### 2.2.2 Covariates

We used soil order as a proxy for forest type. Although it would have been preferable to use soil suborder, we did not have sufficient representation of each across the 43 sites. We only used soil orders that were represented at over 80% of the sites; Ultisols / Inceptisols, Ultisols, Inceptisols, Entisols / Inceptisols, and Entisols. We calculated the proportion of each soil order in the area around each site (1, 5 and 10km radius). The soil orders were grouped and treated as one covariate, so the model either contained all soil covariates or none. Tree cover and land covered by urbanisation proportion of the area around each site covered, calculated from land use/land cover (LCLU) data at 30m resolution (Karra *et al*., 2021) (Table 1). All the covariates were measured within 1km, 5km and 10km grid squares around the central ARU location. Scale and centre transformation was used on all covariates prior to modelling.

**Table 1.**
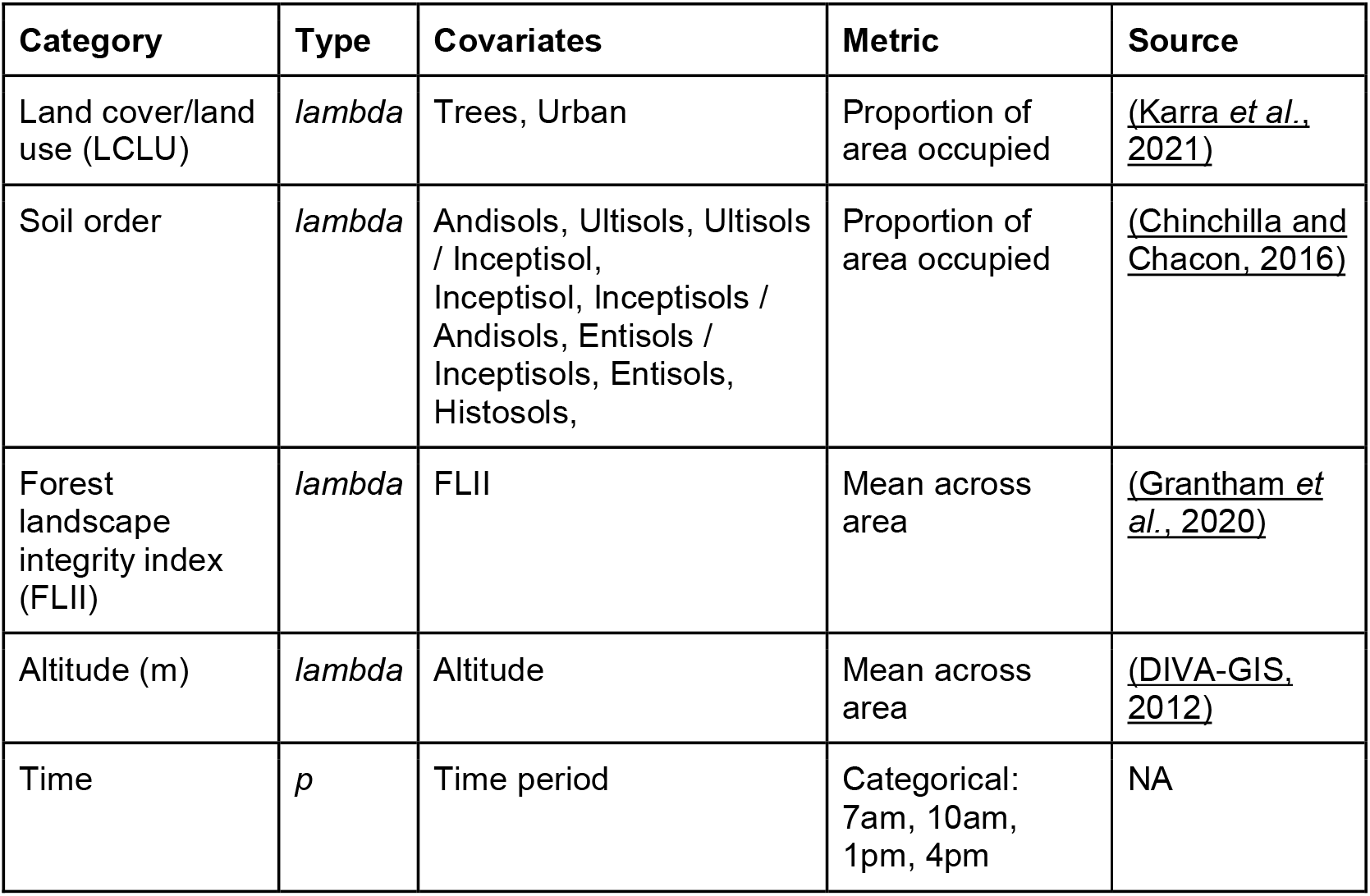
Covariates used in Royle-Nichols model, their definition and source.

#### 2.2.3 Occupancy Modelling

We used the package unmarked (Fiske & Chandler, 2011) using R 4.1.0 (R Core Team 2021) to fit RN models (function *occuRN*). We used a three-step process of model selection. Step 1: we selected the covariates for the detection formula. We evaluated time of day as a continuous covariate from seven to sixteen, time of day as a factor with four levels, date as a number between one and fifty-two, date as a factor with fifty-two levels, and all pairwise combinations of these time and date covariates. Step 2: We evaluated which grid size was the best by comparing the most extensive models at each radius (1, 5 and 10 km). The top-performing radius was then used for the next step. Step 3: every combination of tree cover, urban cover, altitude, forest landscape integrity index and soil type were run, giving a total of 32 models assessed.

For all steps, AIC was used to rank all models; those models within 2 AIC of the top-performing model were considered equal (Burnham & Anderson, 1998). Goodness-of-fit of the top models was assessed using parametric bootstrapping and the sum of squared residuals to calculate fit statistics. The best model fit is one with the lowest difference between the model sum of squared residuals and the mean bootstrapped sum of squared residuals. The model with the best goodness-of-fit was selected to use to estimate abundance. Model coefficients and standard errors (SEs) were calculated on transformed and untransformed covariates.

We used the top model to estimate the number of GGMs at three different levels. The first was abundance at each ARU site, giving a total population estimate for the study area; this was done using the posterior mean of the conditional abundance distribution at each site and estimated using empirical Bayes. Secondly, we extracted all model covariates across the whole of the GGM’s historic range in Costa Rica. We used these and the *predict* function to estimate the potential current population size in Costa Rica. Lastly, we used the ebird range map, smoothed at 9 km resolution (herein “ebird range” - Fink *et al*., 2021), to restrict our population estimate of the current breeding season population in Costa Rica. We used the ebird range to restrict our estimate because our study did not cover the full extent of Costa Rica, so it is difficult to determine where the species is currently extant.

There are five areas where GGMs have been repeatedly reported but are isolated from other GGM areas within Costa Rica and are not included in the smoothed ebird range (Fink *et al*., 2021). These could be remnant breeding populations or seasonal areas. Therefore, we investigated how these isolated areas (herein, “ebird isolated areas”) correspond to our model predictions. If these ebird-isolated areas are predicted to have GGM populations during breeding, this might suggest these are remnant breeding populations. However, we did not include these estimates in our total population estimate as we wanted to develop a conservative population estimate for Costa Rica.

## 3 Results

During the 52 survey days across the 43 sites, GGMs were detected in 592/10712 (5.5%) survey periods at 34 sites (Table S1). Time of day as a factor was the top-performing detection covariate (Table S2), and 10 km was the top-performing grid size (Table S3). There were two models within 2 AIC of each other; one contained all the covariates, and the second did not include the proportion of area covered in urbanisation (Table 2). Parametric bootstrap goodness-of-fit (GOF) tests based on a sum of squared residuals indicated the model without the proportion of area covered in urbanisation fitted the data best, with a difference of 15.88 (model SSE = 466.38, bootstrapped SSE = 482.26, Fig. S1). The only covariate to have a positive model coefficient is tree cover (Fig. 2). Abundance across the ARU sites is estimated to be 208±30 SE, ranging between 0 and 41 individuals per 100km^2^ site (Table S1). The site with the highest abundance is site 38, on the border with Nicaragua, which is Ultisol soil type and has a high proportion of tree cover. There are thirteen sites with an estimate of <1; most are in the southern sites (Fig. 3). These sites are in regions of high agricultural activity associated with low tree cover. There is general agreement between empirical Bayes estimates and predicted abundance across the sites, although some discrepancies might suggest the presence of unmodeled covariates (Fig. S3).

**Table 2.**
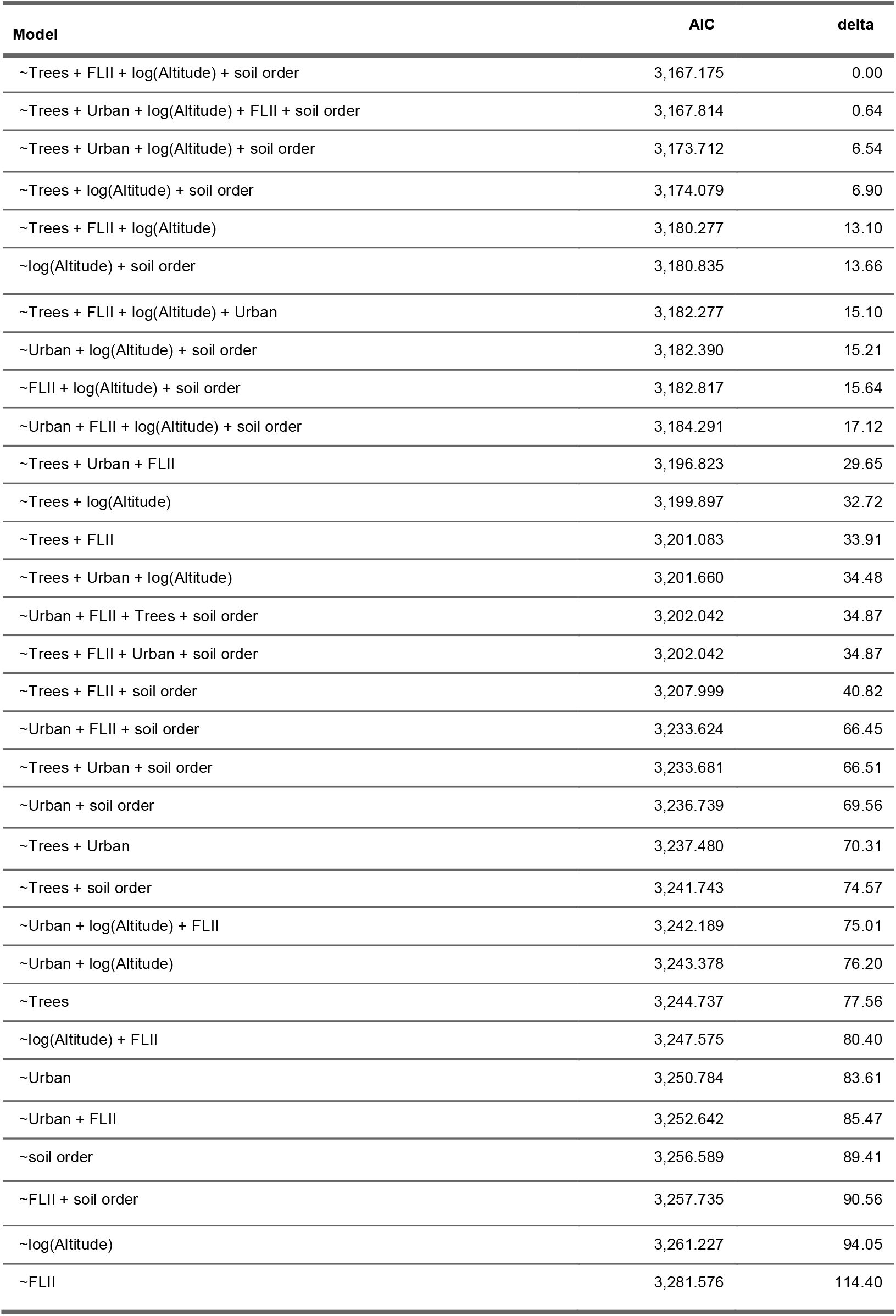
Models, their AIC and delta AIC. The top two models are within 0.64 AIC of each other.

**Figure 2:**
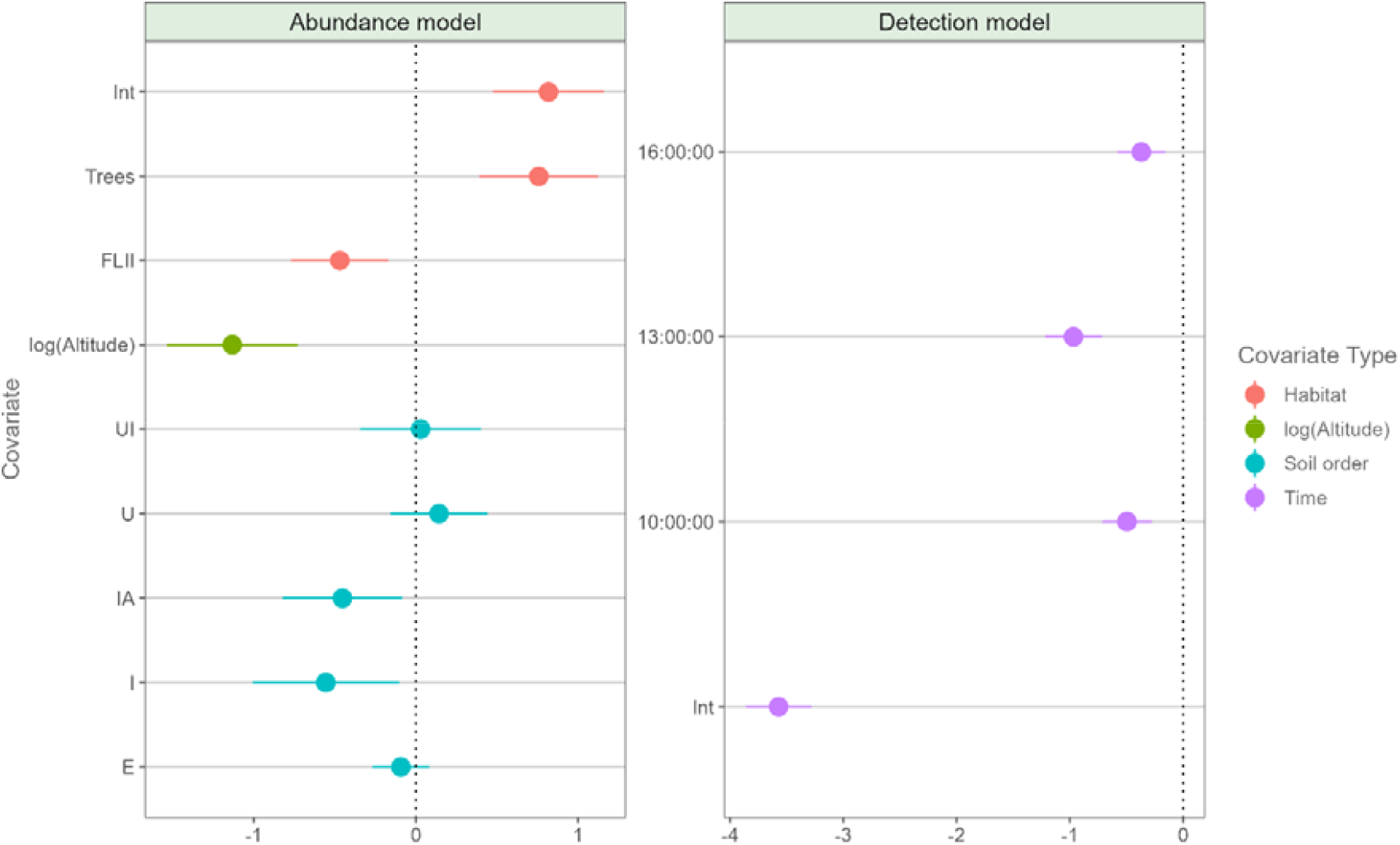
Untransformed model coefficients. The only covariate with a positive effect is Tree cover, whereas all others are either have no significant effect (Ultisol, Ultisol/Inceptisol and Entisol) or negatively affect abundance (FLII, log(Altitude), Inceptisol/Andisol and Inceptisol.

**Figure 3:**
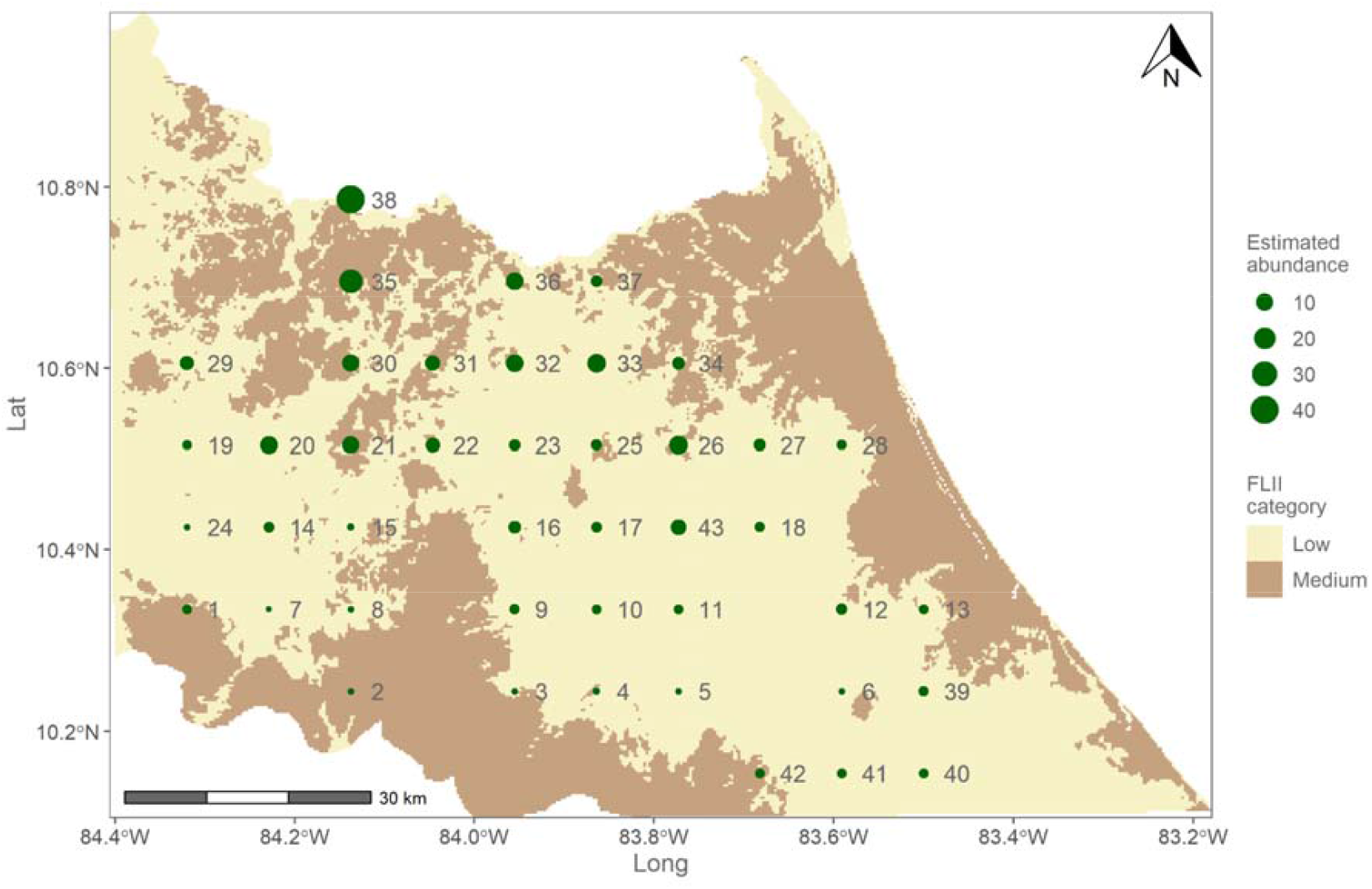
Estimated breeding season abundance across the 43 sites (site ID to the right of locations) shown against forest landscape integrity index (FLII). Higher abundances are seen in the northern areas, compared to the south. FLII has been categorised as per Grantham et al., (2020) into low (0-6), medium (6-9.6) and high (>9.6) integrity.

The RN model predicts a potential breeding population size of 883.09 ± 128 SE throughout the historical range of the GGM, which encompasses the whole Caribbean slope of Costa Rica under 1500m above sea level (a.s.l.) under current environmental conditions. The estimated breeding population size within the ebird range is 485.65±61 SE.

Three of the five ebird isolated areas correspond to areas in which our model predicts there could be significant numbers of GGMs (Fig. 4). The five areas correspond to a potential 49.34 ± 11 SE GGMs; we do not include these in our estimate as we want to be conservative.

**Figure 1:**
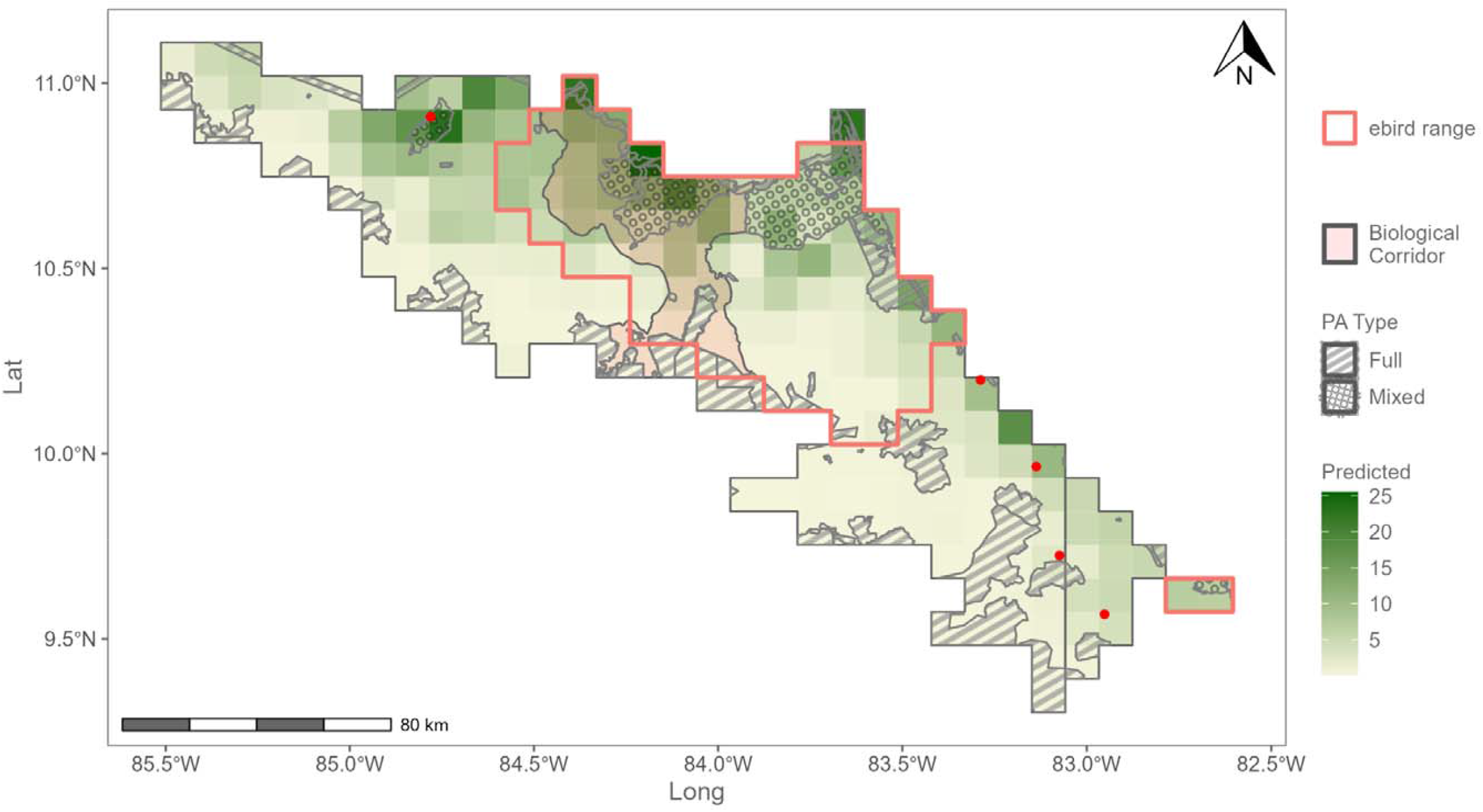
GGM abundance during the peak of the breeding season (Jan - Mar) is largely restricted to lowland areas in the north of Costa Rica on the border with Nicaragua. Significant numbers can be found in the Maquenue Mixed use reserve within the San Juan La Selva biological corridor. Predicted potential breeding season abundance across the historic range of the great green macaw in Costa Rica. Outlined is the Costa Rican range of the great green macaw using ebird data produced by Fink et al., (2021). The biological corridor is the San Juan La Selva biological corridor. Protected area (PA) type “full” is a national park, whereas “mixed” is a mixed-use reserve where forestry and agriculture are permitted under specified limits. Red dots are the five isolated ebird areas.

## 4 Discussion

Robust methods are needed to help increase the effectiveness of species conservation in tropical ecosystems. Here we demonstrate that new monitoring approaches combined with existing statistical methods offer a way to overcome these issues and to do so affordably and at scale. Employing PAM and RN modelling, we estimate the breeding season population of GGM in Costa Rica is 485.65±61SE, which suggests that the global total likely exceeds 500 mature individuals.

Our estimate is similar to a recent non-breeding roost counts survey carried out in Costa Rica that estimated the population to be 336.5 individuals (MRN, 2021). However, as with this current study this will include juveniles who fledged in that year and others not yet considered mature. Tella et al., (2013) and Pacífico et al., (2014) reported that only around 20% of the population of two threatened macaw species were reproductively active in a single season. It is unclear how this equates to the number of mature individuals, as individuals might not be reproductively active in any given year due to their age (Negro 2011), nest-site availability (Cockle et al. 2010) or previous year’s breeding (Berkunsky et al. 2014). Another study estimated that 50% of a blue-throated macaw population was reproductively active over five years (Berkunsky et al. 2014). The Birdlife assessment of the GGM used a figure of 65% to calculate mature population size from total population estimates (BirdLife International 2020), although it is unclear what this is based on. If we use 20% as a lower bound and 65% as an upper bound, the Costa Rican population would consist of 98-315 mature individuals or 49-157 breeding pairs. Considering the Macaw Recovery Network monitored over 40 active nest sites in 2020 within a section of the San Juan La Selva biological corridor (Macaw Recovery Network, 2020), we believe 49 pairs would be a significant underestimate of the Costa Rican population.

Across the ARU sites, our model estimates a population of 206±30SE GGMs. Raw detections and estimates of abundance suggest a hitherto unknown GGM breeding area in the vicinity of sites 26 and 43 (10). To date, the most easterly active nest site was recorded in the vicinity of site 32 (Macaw Recovery Network, *unpub data*); therefore, the high estimates at site 33, 26 and 43 suggests that the breeding area of the GGM in Costa Rica is at least a third larger than previously known. There were also detections made in the group of sites to the southeast of the study area. Although only a small number of detections were made, with a correspondingly low model estimate of individuals (Table S3.1), this suggests a small number of GGMs present in this area during the breeding season. These results show how PAM can identify previously unrecorded areas of occupancy and start to fill the gaps in our knowledge surrounding species distribution. Finally, we have shown that a potential GGM breeding area is currently not being monitored. This means the status and threats to this sub-population are unknown. Site 43 is in an area of high agricultural productivity, with limited accessibility, low income and poor living conditions; this is a concern because these conditions are known to contribute to an increased risk of poaching (UNODC 2012; Pires 2012).

### 4.1 Key Covariates of GGM Abundance

Soil order, our proxy for forest composition, was important for predicting GGM abundance at the regional scale. Although soil order is an essential determining factor of forest community composition (Sollins 1998), other factors such as logging and agricultural intensity influence current forest composition. Two soil orders, Inceptisol and Inceptisol / Andisol, were negatively correlated with GGM abundance. These soil orders are closely related and associated with large river valleys and shore deposits. These soil orders are often found in lowland areas and are often highly anthropogenically altered due to their higher fertility (Eswaran et al. 2002), which is likely why they are negatively correlated with GGM abundance. The other soil orders, Entisol, Ultisol and Ultisol / Inceptisol, do not have a significant effect on GGM abundance. For Ultisol and Ultisol / Inceptisol soil types, this was not expected, because these soil types are associated with higher mountain almond density, a key breeding season resource of the GGMs. Mountain almond grows well in alluvial or sandy soils with an acidic profile (Flores 1992; Vidal-Riveros 2004) which is not the case for Entisol, Inceptisol or Inceptisol / Andisol orders (Chun, 2008). The soil suborder, Humults, which is a Ultisol soil, is a key predictor of mountain almond tree density in northern Costa Rica (Chun 2008). Ultisol soils are formed from a process of clay mineral weathering of old land surfaces, have a lower pH and have subsurface horizons of illuvial clay accumulations (Brady & Weil, 1996). Therefore our finding that Ultisol soil order is not positively associated with GGM abundance goes against the hypothesis that the distribution of mountain almonds is a limiting factor of GGM breeding season abundance in Costa Rica.

Sites with higher abundance were generally found further north, in areas of low altitude and higher forest cover. These areas are more suited to mountain almond growth (Flores 1992; Chun 2008) and are also less developed and have had some form of protection since the 1990s, allowing for some natural regeneration (Fagan et al. 2013). Northern sites are also closer to the Indio-Maiz biological reserve in Nicaragua, where there are large tracts of primary forest with lower relative human disturbance (Gamboa and Sofía, 2021). The abundance of GGMs in the Indio-Maiz biological reserve is uncertain. A small section on the western edge of the reserve was surveyed in 2008, and the population was estimated to be 303 ± 514 individuals (Monge et al., 2010). Although the methodology used was not robust, comparing the raw number of GGMs sighted in each area suggests that this area has more GGM activity than the Costa Rica sites monitored during the 2008 study. High abundance at the sites in the north is likely due to the amount of protection, poor accessibility and vicinity to the Indio-Maiz biological reserve.

The Costa Rican Caribbean lowland rainforest (below 500m a.s.l.) is a degraded ecosystem, with 74% categorised as having low FLII (Fig 3; Grantham *et al*., 2020); this means that it is highly anthropogenically disturbed. In areas of high anthropogenic disturbance, we find fewer mature hardwood trees (DeWalt et al. 2003), which are those that support GGMs with food and nesting resources (Powell et al. 1999; Monge et al. 2003). So, perhaps counter-intuitively, the forest landscape integrity index (FLII) has a small negative effect on abundance. This means that more intact forest is weakly negatively associated with lower GGM abundance. In Costa Rica, the GGM is heavily reliant on the mountain almond as a nesting and food resource; ∼85% of their nests are located in this species, and up to 90% of their diet consists of mountain almond nuts during the breeding season (Monge *et al*., 2003, 2012; Chassot, Arias and Powell, 2007; Lewis *et al*., *in prep*.). Mountain almond is a valuable hardwood prized for its ability to resist rot and termites (Flores 1992). Since Costa Rica has experienced one of the highest deforestation rates globally (de Camino et al. 2000), this vital resource was impacted by timber extraction. However, large-scale exploitation was limited until the 1980s, when improved saws and milling technology were developed to cope with mountain almonds’ density and crystalline deposit content (Flores 1992; Butterfield 1995). The mountain almond was regionally protected in 1999 and nationally protected in 2008 (Powell et al. 1999; Sancho 2008). Consequently, mountain almond trees are found in relatively high densities in some forested areas and scattered throughout low integrity areas such as pastures and secondary forests, especially throughout the eastern half of the San Juan La Selva biological corridor (Chun, 2008; Macaw Recovery Network, *unpub. data*). These remnant mountain almond trees may allow GGMs to utilise low integrity, secondary forest and otherwise barren pasturelands.

Discrepancies exist between the ebird range and our model in central areas of the historical range (Fig. 4) because the ebird range is a year-round occupancy range. In contrast, we are estimating the population during only the breeding season. For example, we know GGMs utilise areas of higher altitude in the south and southwest of the ebird range during the non-breeding season (Powell et al. 1999; Monge et al. 2003; Macaw Recovery Network 2021). This explains the lack of predicted abundance in these areas from our model. Our predictions of high abundance areas in the north agree with what we see on the ground (TL *pers. obs*) and ebird data (Fink et al. 2021). Another area where there is a lack of predicted GGMs is over areas of expansive monoculture crops such as banana and pineapple in the southeast of the ebird range; this is likely because the ebird range is the extent of the range and does not have detail within it.

The isolated population identified by Fink *et al*. (2021) in the northwest, where our model predicts a significant population, corresponds with Refugio de Vida Silvestre Caño Negro. This is a protected area with higher tree cover than non-protected areas. Areas to the country’s southeast are predicted to have low but significant numbers of birds. This is interesting as the far southeast is the location of a reintroduction of GGM in 2011. Before the reintroduction, there were only a few sporadic sights of GGMs in the region (ebird 2021).

### 4.2 Potential biases and limitations

The historical range of the GGM in Costa Rica extends throughout the Caribbean slope of Costa Rica to a maximum height of 1500m a.s.l. (Powell et al. 1999; Monge et al. 2003). Our model predicts that there could be a population of 883±128 SE in this area under current environmental conditions, whereas within the ebird range, there is an estimated 485±65 SE. The discrepancies between our estimates are due to the northwest and the east of the historical range. This suggests that we lack information in our covariates. We assumed that the soil type represents the current forest type, whereas it represents the forest type if the forest is primary. This is particularly important when considering the close relationship between mountain almonds and breeding for the GGM. For example, in Costa Rica, the initial ban on mountain almond extraction only applied to the area between the San Juan, San Carlos, and Sarapiqui rivers (Powell *et al*., 1999) in the eastern half of the San Juan La Selva biological corridor (Fig. 4). Chun (2008) suggests that mountain almond distribution in the rest of the San Juan La Selva biological corridor is lower than expected, given environmental conditions. This pattern will likely be the same in areas outside the San Juan La Selva biological corridor open to exploitation until 2008 (Sancho 2008). Establishing the current distribution and density of mountain almonds across the Caribbean lowland would surely provide important information that could be used to aid the conservation of the GGM.

RN models assume that the target populations are closed. We attempted to limit the probability that this assumption would be violated by placing ARUs 10 km apart. However, if a large proportion of the population is composed of non-breeding individuals located outside breeding sites, this poorly observed fraction of the population will affect population estimates in ways that are challenging to quantify (Penteriani et al., 2011; Negro 2011). For example, field observations suggest that juveniles may follow their parents for up to 3 years (Jimenez *pers comms*.; TL *pers obs*), but once juveniles are independent, their behaviour is unclear. These individuals may spend their time in locations with high-value food species, such as mountain almonds. However, the extent to which these areas overlap with breeding sites is unknown.

### 4.3 Conclusion

Challenges with estimating abundance for highly mobile species such as parrots are well documented (Casagrande & Beissinger, 1997; Dénes, Tella and Beissinger, 2018; Tella *et al*., 2021). However, there have been few previous attempts to estimate the Costa Rican GGM population. Powell *et al*., (1999) estimated that the Costa Rican GGM population was around 210 individuals in the 1990s. A subsequent population census estimated the population to be 303 ± 514 (Monge et al. 2010). The wide confidence intervals of Monge *et al*., (2010) demonstrate how challenging obtaining robust estimates of parrot abundance is. We have shown that PAM and RN modelling can be used together to provide robust abundance information for critically endangered parrot species such as the GGM. Our model highlighted previously unknown breeding areas, enabled us to resolve breeding habitat associations for the GGM in Costa Rica and allowed us to predict potential GGM distribution. We have demonstrated at a population scale that these tools offer conservationists an efficient and effective way to understand population dynamics. With a high proportion of parrot species threatened globally (IUCN 2020b), this tool will be essential in conservation planning and evaluation. PAM technology has enormous potential to facilitate such assessments because it is easily scalable, recordings can be stored and re-analysed as machine learning, and abundance estimation techniques become more advanced.

---

**Figure 1:**
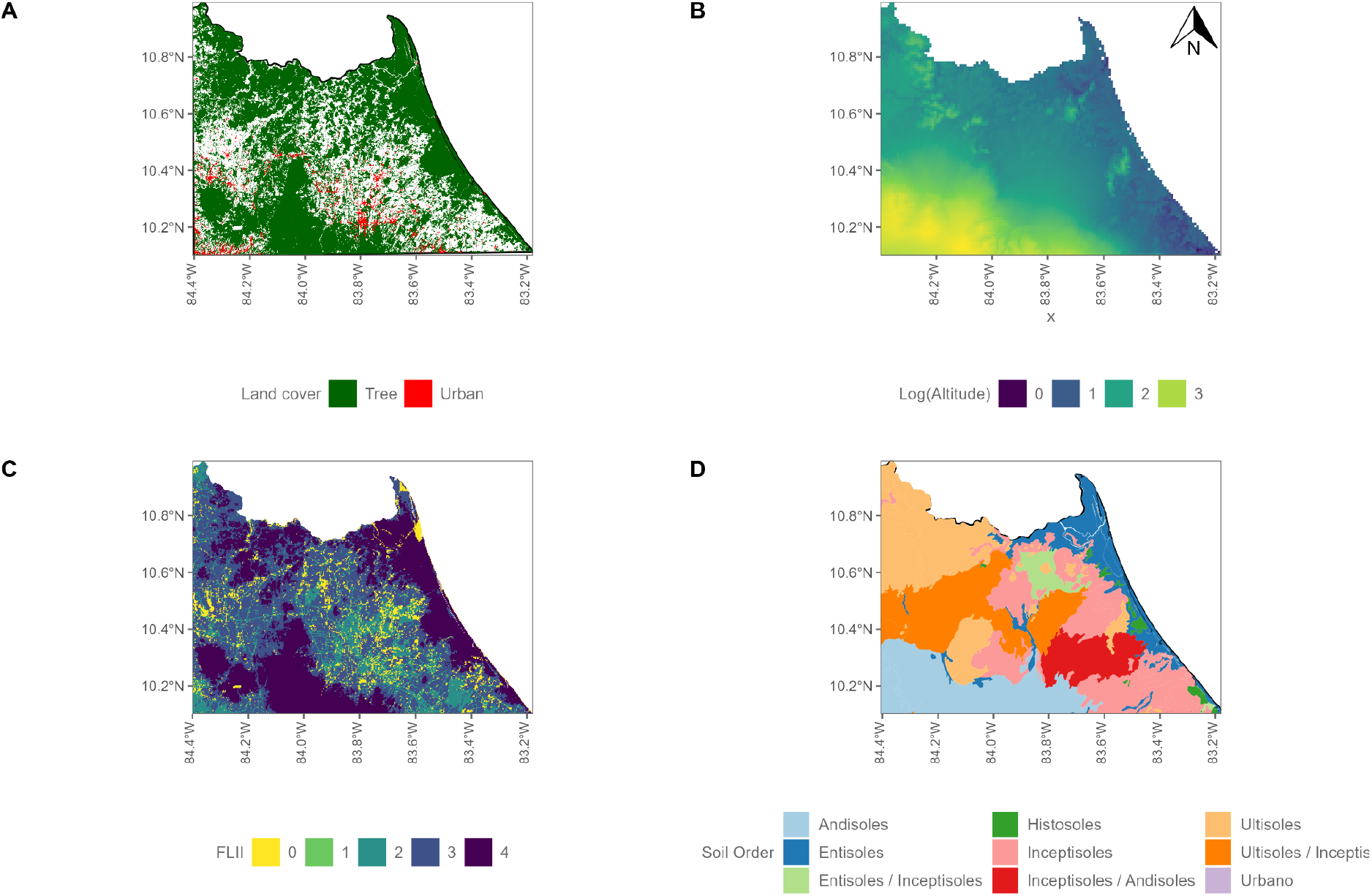
Model covariates across the study site with ARU locations A) Land cover type. B) Altitude (m) C) Forest Landscape Integrity Index (FLII) scaled and centred, higher values indicate more intact forest. D) Soil order (Chinchilla and Chacon, 2016).

**Table 1:**
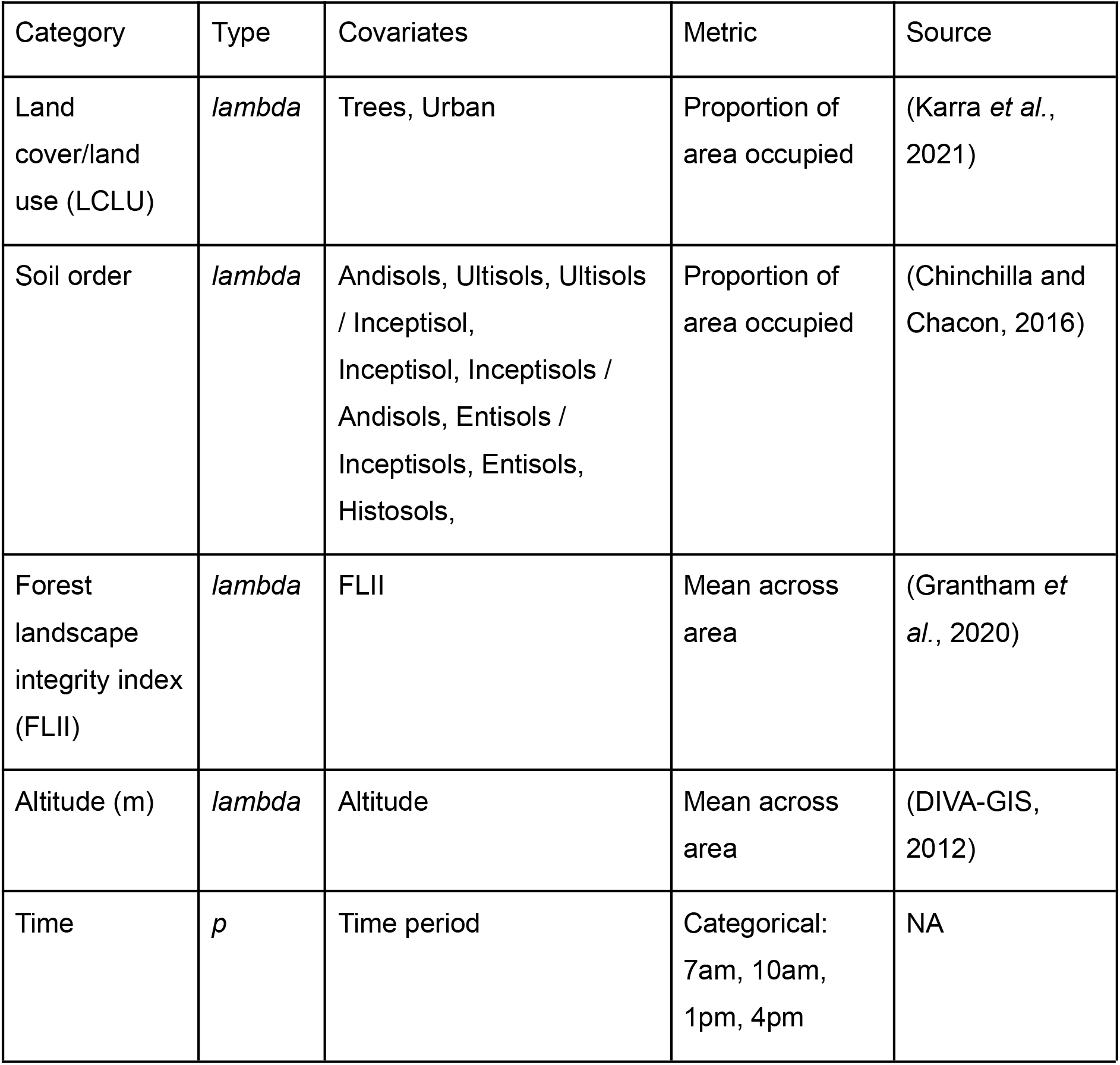
Definitions and source of all covariates used in Royle-Nichol models

**Table 2:**
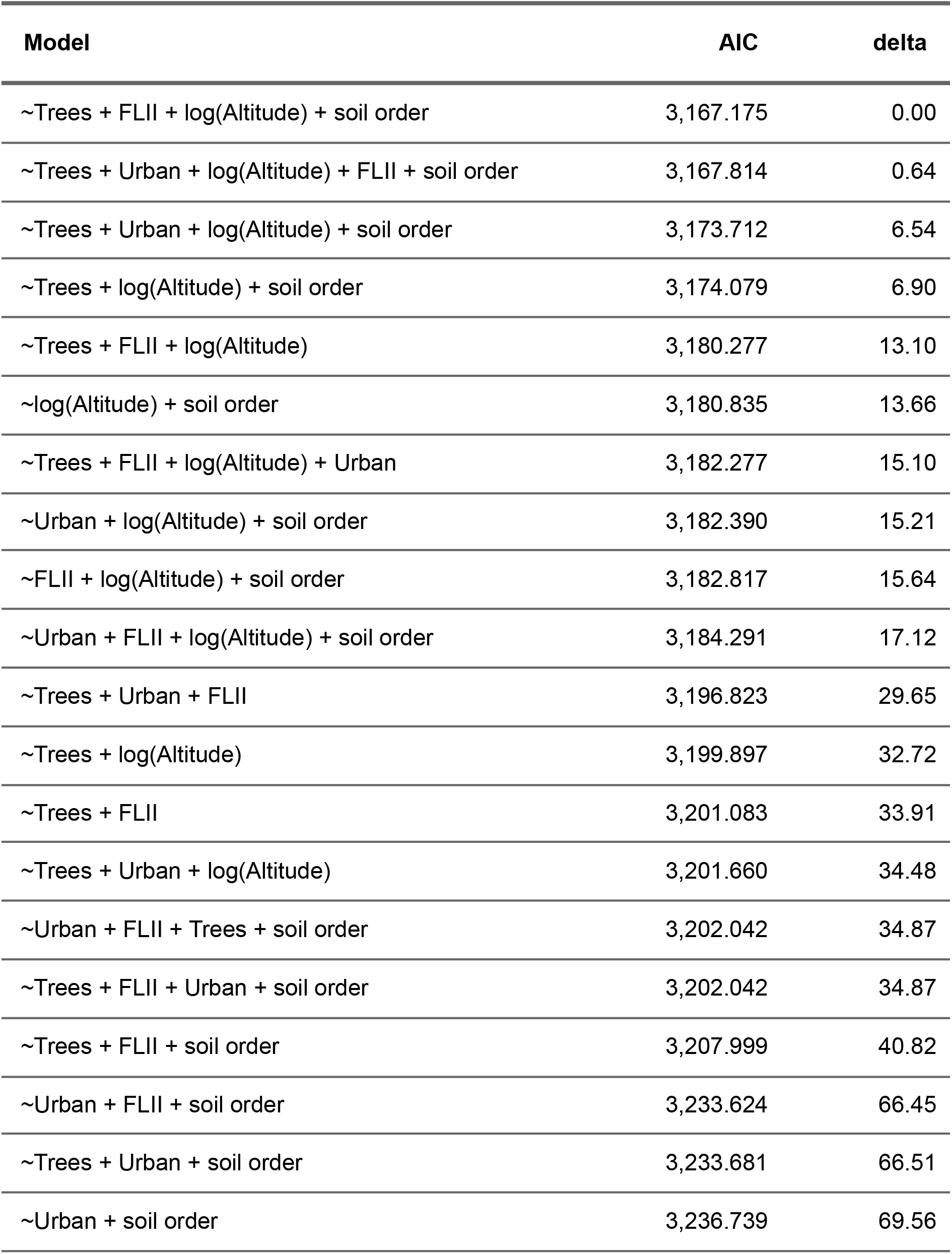
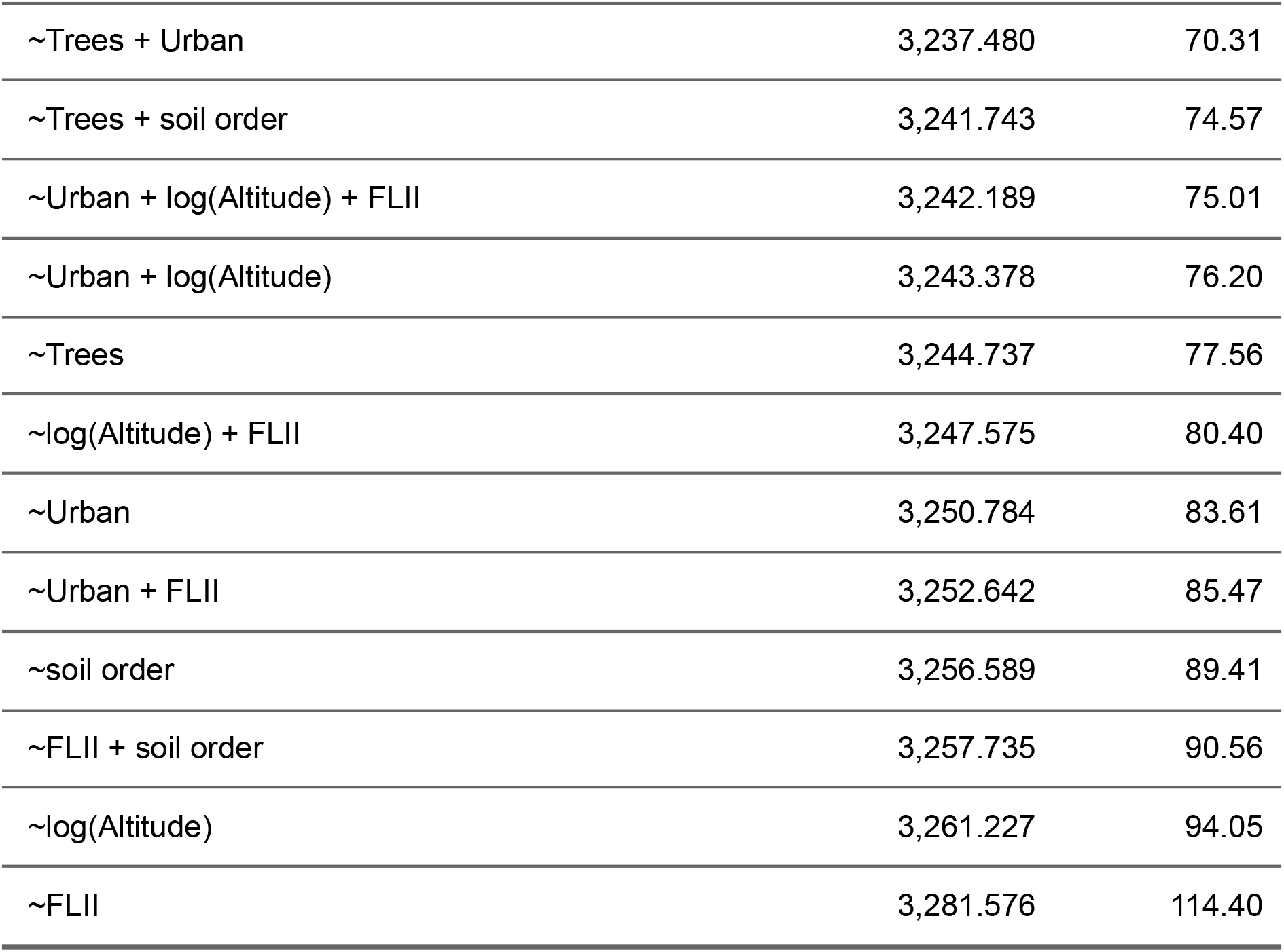
Model AIC table and delta weights. The top two models, within delta 2AIC of the top model, were the global model and the global model without urban area cover.

**Figure 2:**
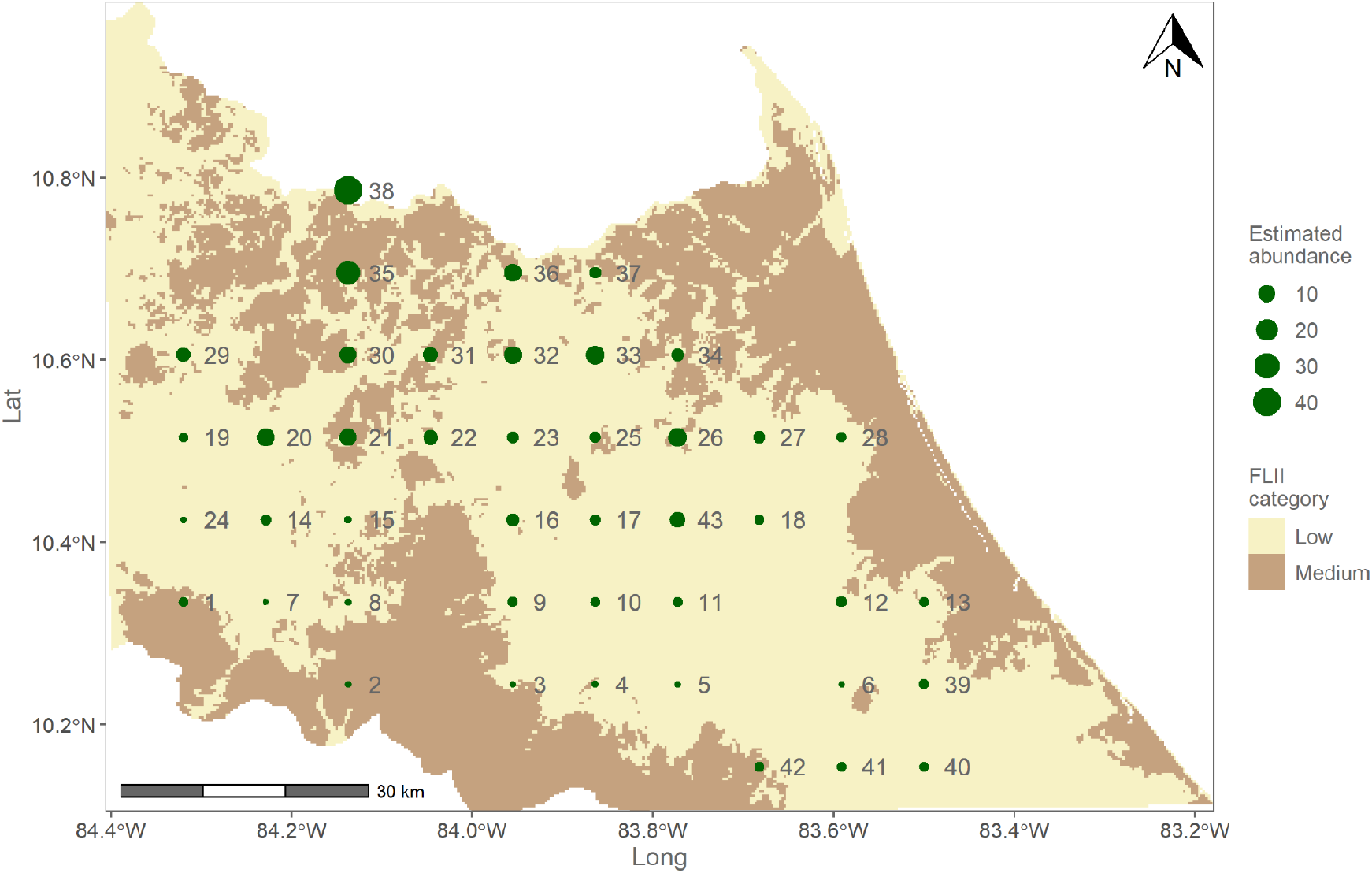
Estimated breeding season abundance across the 43 sites (site ID to the right of locations) shown against forest landscape integrity index (FLII). Higher abundances are seen in the northern areas, compared to the south. FLII has been categorised as per Grantham *et al*., (2020) into low (0-6), medium (6-9.6) and high (>9.6) integrity.

**Figure 3:**
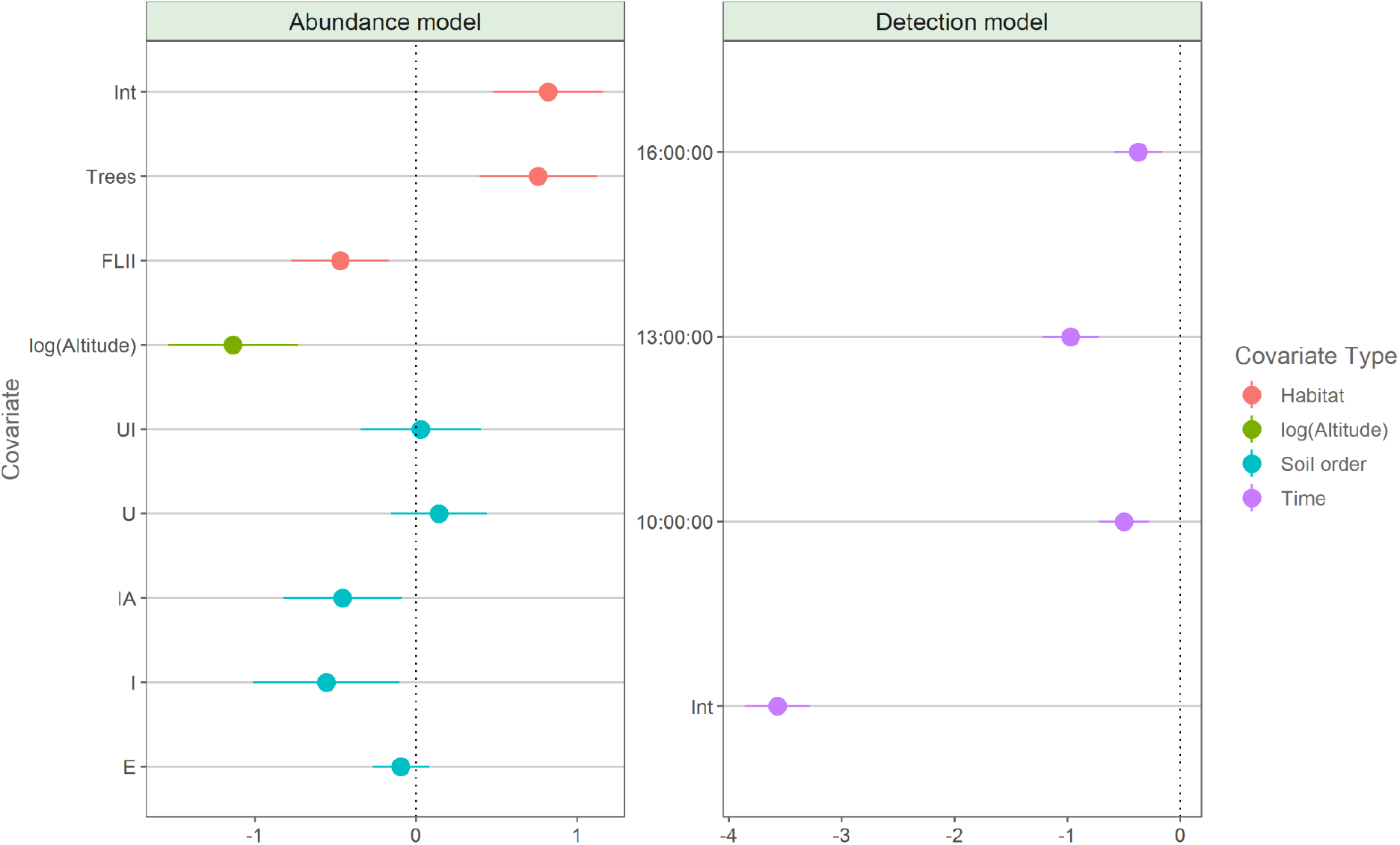
Model coefficients and standard errors (SE) from the top model. Abundance (*lam*) and detection (*p*) covariates. The intercept in *p* represents the 7:00:00 survey period. EI = soil order Entisols / Inceptisols, UI = soil order Ultisols / Inceptisols, U = soil order Ultisols, I = Inceptisols, E = Entisols.

**Figure 4:**
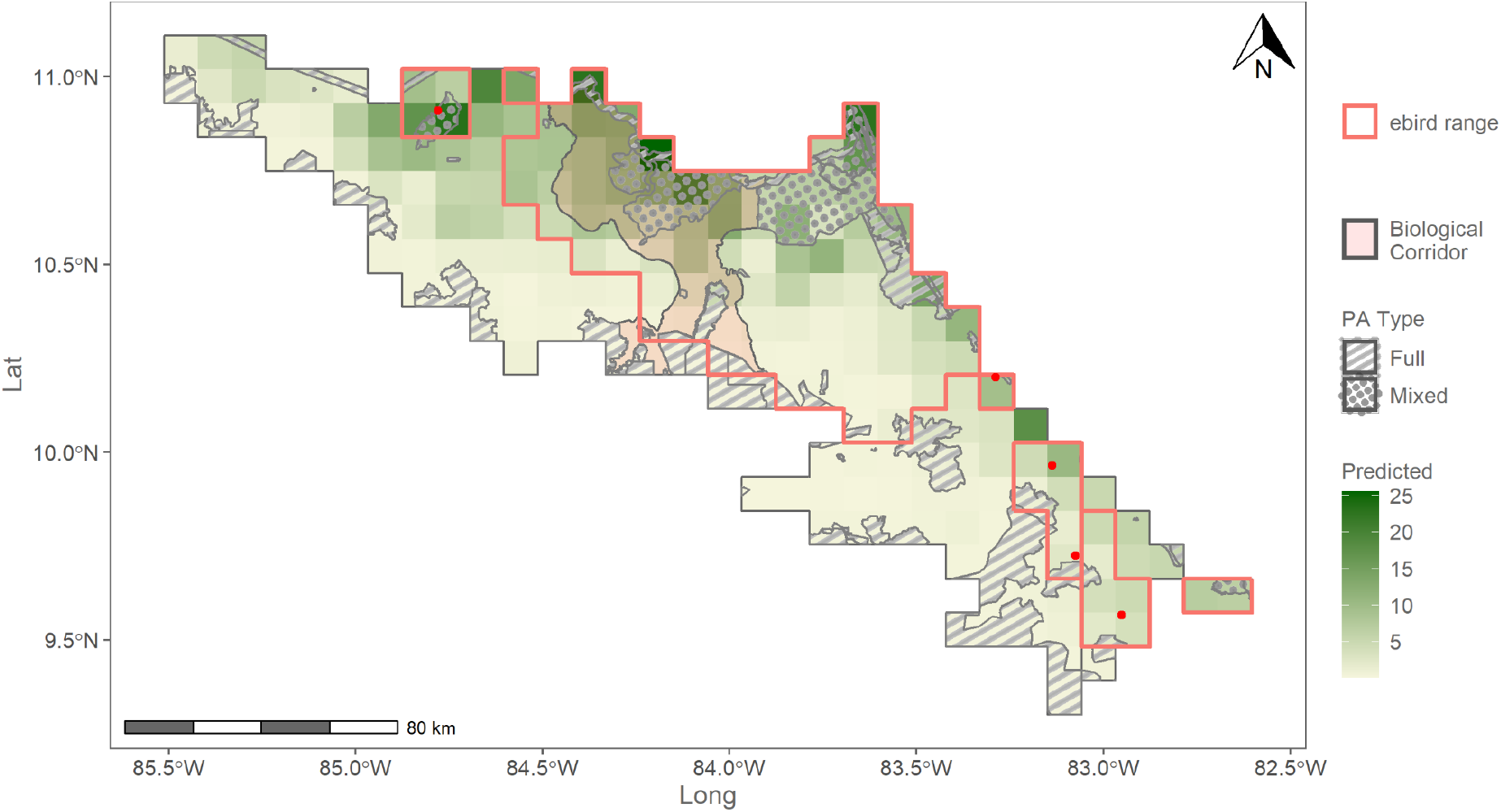
GGM abundance during the peak of the breeding season (Jan - Mar) is largely restricted to lowland areas in the north of Costa Rica on the border with Nicaragua. Significant numbers can be found in the Maquenue Mixed use reserve within the San Juan La Selva biological corridor. Predicted potential breeding season abundance across the historic range of the great green macaw in Costa Rica. Outlined is the Costa Rican range of the great green macaw using ebird data produced by Fink *et al*., (2021). The biological corridor is the San Juan La Selva biological corridor. Park type “full” is a national park, whereas “mixed” is a mixed-use reserve where forestry and agriculture are permitted under specified limits.

