## Supplementary Materials 1 for "Using passive acoustic monitoring to estimate the abundance of a critically endangered parrot, the great green macaw (*Ara ambiguus*)": Supplementary Materials 1.html

RN code


### RN code

#### TL

#### 2022-11-28

### The Setup

#### Packages

First to install the packages needed for this script:

```
library(unmarked)
library(tidyverse)
library(lubridate)
library(scatterpie)
library(GGally)
library(sf)
library(fs)
library(ggpubr)
library(ggspatial)
library(raster)
library(ggtext)
library(emdbook)
library(ggnewscale)
library(ggpattern)
library(knitr)
library(expss)
library(flextable)
```

#### Load data

```
data_RN <- read_csv("./Data/Data_RN.csv")
```

#### Detection covariates

There are a number of different temporal detection covariates that
could be used. So lets test each one and their combinations and see
which is the best. The top detection covariate(s) will then be used in
all subsequent models.

```
# load time covariates
cov_time <- read_csv("./Data/time_covariates.csv")

#  time as a factor
cov_time_factor <- cov_time %>% 
  dplyr::select(site, datetime, time) %>%  
  mutate(time = as.factor(time)) %>% 
  pivot_wider(names_from = "datetime", values_from = "time", names_sort = T)

#  time as a number 
cov_time_numeric <- cov_time %>% 
  dplyr::select(site, datetime, hour) %>%  
  pivot_wider(names_from = "datetime", values_from = "hour", names_sort = T)

# date covariates
cov_date <- cov_time %>% 
  dplyr::select(site, datetime) %>%
  mutate(date = as.factor(as.Date(datetime))) %>% 
  pivot_wider(names_from = "datetime", values_from = "date", names_sort = T) 

# create a list of covariate
cov_list <- list(cov_time_factor, cov_time_numeric, cov_date) %>% 
  map(~dplyr::select(., -site)) %>% 
  set_names(nm = c("time.of.day.factor","time.of.day.numeric","date.factor"))
```

In summary, variable definitions: `time.of.day.factor` =
time of day as a factor `time.of.day.numeric` = time of day
as a number ``date.factor``` = date as a factor

##### Site covariates

For this modelling I am using five basic site level covariates:

- Mean log\_Altitude of site
- Tree cover
- Urban cover
- Forest Landscape Integrity index (FLII)
- Soil order

See [Covariate extraction] for more information on these and how they
were extracted

##### Load covariates

```
site_covs <- as_tibble(dir_ls("./Data/", regexp = "covariate_site_grid_extended")) %>% 
  mutate(width = str_extract(as.character(value), "[0-9]+")) %>% 
  group_split(width) %>% 
  map(~read_csv(.$value) %>% 
        dplyr::select(-.id) %>% 
        mutate(log_Altitude = log10(Altitude)))

width <- c("1000", "10000", "5000")

site_covs %>% 
  map(~dplyr::select(., Trees, Urban, FLII, log_Altitude, ULTISOLES, ULTISOLES_INCEPTISOLES, ENTISOLES, 
              ENTISOLES_INCEPTISOLES,  INCEPTISOLES,  INCEPTISOLES_ANDISOLES) %>% 
        mutate(id = row_number()) %>% 
        pivot_longer(cols = Trees:INCEPTISOLES_ANDISOLES, names_to = "Covariate", values_to = "values") %>% 
        mutate(values = as.factor(values)) %>% 
        group_by(Covariate, values) %>% 
        tally() %>% 
        filter(values == 0 & n >= 35)) %>% 
  map2(., width, ~.x %>% mutate(width = .y)) %>% 
  bind_rows() %>% 
  arrange(as.numeric(width))
```

```
## # A tibble: 6 x 4
## # Groups:   Covariate [3]
##   Covariate              values     n width
##   <chr>                  <fct>  <int> <chr>
## 1 ENTISOLES              0         40 1000 
## 2 ENTISOLES_INCEPTISOLES 0         41 1000 
## 3 INCEPTISOLES_ANDISOLES 0         37 1000 
## 4 ENTISOLES_INCEPTISOLES 0         38 5000 
## 5 INCEPTISOLES_ANDISOLES 0         37 5000 
## 6 ENTISOLES_INCEPTISOLES 0         35 10000
```

```
# now to scale and center everything
site_covs <- site_covs %>% 
  map(~mutate(., across(everything(), ~scale(.x, center = T))))
```

Lets check if there are any correlations between the numeric
covariates

```
# lets check correlation between covariates

site_plot <- as_tibble(dir_ls("./Data/", regexp = "covariate_site_grid_extended")) %>% 
  mutate(width = str_extract(as.character(value), "[0-9]+")) %>% 
  filter(width == 10000) %>%
  group_split(width) %>% 
  map(~read_csv(.$value)) %>%
  bind_rows() %>% 
  mutate(log_Altitude = log10(Altitude)) %>% 
  dplyr::select(., .id, Trees, Urban, log_Altitude, FLII, ULTISOLES, INCEPTISOLES, ENTISOLES,
                     ENTISOLES_INCEPTISOLES, INCEPTISOLES_ANDISOLES) %>% 
  pivot_longer(cols = 2:ncol(.), names_to = "Varibles", values_to = "values") %>% 
  mutate(covariate = str_to_title(gsub("_", "/\n", Varibles, fixed=TRUE)),
         covariate = ifelse(covariate == "Flii", "FLII", covariate),
         covariate = ifelse(covariate == "Log/\nAltitude", "log(Altitude)", covariate),
         covariate = str_replace_all(covariate, "soles", "sols")) %>% 
  dplyr::select(-Varibles) %>% 
  pivot_wider(names_from = "covariate", values_from = "values") %>% 
  dplyr::select(-.id)


ggpairs(data = site_plot) +
      theme(panel.grid.major = element_blank(),
          panel.grid.minor = element_blank(),
          panel.background = element_blank(),
          panel.border = element_rect(fill = NA, colour = "gray40"),
          axis.ticks = element_line(colour = "gray40"),
          axis.title = element_text(size = 10, colour = "gray40"),
          axis.text = element_text(size = 10),
          plot.margin = margin(5,5,5,5, unit = "pt"),
          strip.background =element_rect(colour = "gray40", fill = "honeydew2"),
          strip.text = element_text(colour = "gray40", size = 10),
          legend.position = "bottom",
          strip.text.y.right = element_text(angle = 0))
```

Trees and FLII are massively correlated, this makes sense as where
there is higher forest cover there is more likely to be higher forest
integrity. I will keep both covariates as they are measuring slightly
different things. We can always drop one later, but for now we can keep
it in to give us a full suite of options.

### Royle-Nichol models

#### Step 1: Detection covariate testing

First we have to define the detection covariate(s) that are going to
be used. TO do this we evaluate all possible detection covariate
combinations and select the lowest AIC

```
# time as a factor only
umf_1 <- unmarkedFrameOccu(y = data_RN, siteCovs = site_covs[[1]], obsCovs = cov_list)

# run occuRN with different combinations:
fit_1 <- occuRN(~1 ~1, data = umf_1, K = 50, engine = "R")
fit_2 <- occuRN(~time.of.day.numeric ~1, data = umf_1, K = 50, engine = "R")
fit_3 <- occuRN(~date.factor ~1, data = umf_1, K = 50, engine = "R")
fit_4 <- occuRN(~time.of.day.factor ~1, data = umf_1, K = 50, engine = "R")
fit_5 <- occuRN(~time.of.day.numeric + date.factor ~1, data = umf_1, K = 50, engine = "R")
fit_6 <- occuRN(~time.of.day.factor + date.factor ~1, data = umf_1, K = 50, engine = "R")

# find the lowest AIC
detection_table <- list(fit_1, fit_2, fit_3, fit_4, fit_5, fit_6) %>% 
  map(~.@AIC) %>% 
  set_names(nm = c("Intercept", "Time of day (numeric)", "Date (factor)", "Time of day (factor)", 
                   "Time of day (numeric) + Date (factor)", "Time of day (factor) + Date (factor)")) %>% 
  bind_rows() %>% 
  pivot_longer(cols = 1:6, names_to = "Model", values_to = "AIC") %>% 
  arrange(AIC) %>% 
  mutate(delta = round(AIC -(AIC[1]),2),
         delta = replace_na(delta, 0.00)) 

# create full model flextable to export to word
det_ft <- flextable(detection_table, 
  col_keys = c("Model", "AIC", "delta")) %>% 
  width(width = c(4,1,1)) %>% 
  padding(padding = 1) %>% 
  theme_vanilla()

det_ft
```

| Model | AIC | delta |
| --- | --- | --- |
| Time of day (factor) | 3,283.727 | 0.00 |
| Time of day (factor) + Date (factor) | 3,316.236 | 32.51 |
| Time of day (numeric) | 3,322.652 | 38.93 |
| Intercept | 3,340.115 | 56.39 |
| Time of day (numeric) + Date (factor) | 3,355.837 | 72.11 |
| Date (factor) | 3,373.232 | 89.51 |

```
# save
save_as_docx(det_ft, path = "./model_table_detection.docx")
```

So the best model of the ones I have run is using time of day as a
factor.

The fact that models with date in them are not within 30 AIC suggests
that there is no change in detection across the period we are looking
at. This is good as the reason we are using it is because we want a
period where birds are active, breeding before chicks fledge.

#### Step 2: Scale

We have to see at which scale to use for the modelling stage. To do
this we create one model for each scale, this model will be the biggest
model possible, using all the covariates. The model with the lowest AIC
will be the scale we use in the next step.

```
umf_2 <- unmarkedFrameOccu(y = data_RN, siteCovs = site_covs[[3]], obsCovs = cov_list)
umf_3 <- unmarkedFrameOccu(y = data_RN, siteCovs = site_covs[[2]], obsCovs = cov_list)

small_1 <- occuRN(~time.of.day.factor ~Trees + Urban + log_Altitude + FLII + ULTISOLES + ULTISOLES_INCEPTISOLES  
                  + INCEPTISOLES, data = umf_1, K = 50, engine = "R")
med_2 <- occuRN(~time.of.day.factor ~Trees + Urban + log_Altitude + FLII + ULTISOLES + ULTISOLES_INCEPTISOLES  
                  + ENTISOLES + INCEPTISOLES, data = umf_2, K = 50, engine = "R")
large_3 <- occuRN(~time.of.day.factor ~Trees + Urban + log_Altitude + FLII + ULTISOLES + ULTISOLES_INCEPTISOLES  
                  + ENTISOLES + INCEPTISOLES + INCEPTISOLES_ANDISOLES, data = umf_3, K = 50, engine = "R")
# find the lowest AIC
table_size <- list(small_1, med_2, large_3) %>% 
  map(~.@AIC) %>% 
  set_names(nm = c("1 km", "5 km", "10 km")) %>% 
  bind_rows() %>% 
  pivot_longer(cols = 1:3, names_to = "Model", values_to = "AIC") %>% 
  arrange(AIC) %>% 
  mutate(delta = round(AIC -(AIC[1]),2),
         delta = replace_na(delta, 0.00)) 

# create full model flextable to export to word
size_ft <- flextable(table_size, 
  col_keys = c("Model", "AIC", "delta"))  %>% 
  width(width = c(4,1,1)) %>% 
  padding(padding = 1) %>% 
  theme_vanilla()

size_ft
```

| Model | AIC | delta |
| --- | --- | --- |
| 10 km | 3,167.814 | 0.00 |
| 5 km | 3,178.637 | 10.82 |
| 1 km | 3,207.277 | 39.46 |

```
# save
save_as_docx(size_ft, path = "./model_table_size.docx")
```

#### Step 3: Abundance covariates

```
# 1 covariate
large_1 <- occuRN(~time.of.day.factor ~Trees, data = umf_3, K = 50, engine = "R")
large_2 <- occuRN(~time.of.day.factor ~Urban, data = umf_3, K = 50, engine = "R")
large_3 <- occuRN(~time.of.day.factor ~log_Altitude, data = umf_3, K = 50, engine = "R")
large_4 <- occuRN(~time.of.day.factor ~FLII, data = umf_3, K = 50, engine = "R")
large_5 <- occuRN(~time.of.day.factor ~ULTISOLES + ULTISOLES_INCEPTISOLES  
                  + ENTISOLES + INCEPTISOLES + INCEPTISOLES_ANDISOLES, data = umf_3, K = 50, engine = "R")

# 2 covariates
large_6 <- occuRN(~time.of.day.factor ~Trees + Urban, data = umf_3, K = 50, engine = "R")
large_7 <- occuRN(~time.of.day.factor ~Trees + log_Altitude, data = umf_3, K = 50, engine = "R")
large_8 <- occuRN(~time.of.day.factor ~Trees + FLII, data = umf_3, K = 50, engine = "R")
large_9 <- occuRN(~time.of.day.factor ~Trees + ULTISOLES + ULTISOLES_INCEPTISOLES  
                  + ENTISOLES + INCEPTISOLES + INCEPTISOLES_ANDISOLES, data = umf_3, K = 50, engine = "R")
large_10 <- occuRN(~time.of.day.factor ~Urban + FLII, data = umf_3, K = 50, engine = "R")
large_11 <- occuRN(~time.of.day.factor ~Urban + log_Altitude, data = umf_3, K = 50, engine = "R")
large_12 <- occuRN(~time.of.day.factor ~Urban + ULTISOLES + ULTISOLES_INCEPTISOLES  
                  + ENTISOLES + INCEPTISOLES + INCEPTISOLES_ANDISOLES, data = umf_3, K = 50, engine = "R")
large_13 <- occuRN(~time.of.day.factor ~log_Altitude + FLII, data = umf_3, K = 50, engine = "R")
large_14 <- occuRN(~time.of.day.factor ~log_Altitude + ULTISOLES + ULTISOLES_INCEPTISOLES  
                  + ENTISOLES + INCEPTISOLES + INCEPTISOLES_ANDISOLES, data = umf_3, K = 50, engine = "R")
large_15 <- occuRN(~time.of.day.factor ~FLII + ULTISOLES + ULTISOLES_INCEPTISOLES  
                  + ENTISOLES + INCEPTISOLES + INCEPTISOLES_ANDISOLES, data = umf_3, K = 50, engine = "R")

# 3 covariates 
large_16 <- occuRN(~time.of.day.factor ~Trees + Urban + log_Altitude, data = umf_3, K = 50, engine = "R")
large_17 <- occuRN(~time.of.day.factor ~Trees + Urban + FLII, data = umf_3, K = 50, engine = "R")
large_18 <- occuRN(~time.of.day.factor ~Trees + Urban + ULTISOLES + ULTISOLES_INCEPTISOLES  
                  + ENTISOLES + INCEPTISOLES + INCEPTISOLES_ANDISOLES, data = umf_3, K = 50, engine = "R")
large_19 <- occuRN(~time.of.day.factor ~Trees + FLII + log_Altitude, data = umf_3, K = 50, engine = "R")
large_20 <- occuRN(~time.of.day.factor ~Trees + FLII + ULTISOLES + ULTISOLES_INCEPTISOLES  
                  + ENTISOLES + INCEPTISOLES + INCEPTISOLES_ANDISOLES, data = umf_3, K = 50, engine = "R")
large_21 <- occuRN(~time.of.day.factor ~Trees + log_Altitude + ULTISOLES + ULTISOLES_INCEPTISOLES  
                  + ENTISOLES + INCEPTISOLES + INCEPTISOLES_ANDISOLES, data = umf_3, K = 50, engine = "R")
large_22 <- occuRN(~time.of.day.factor ~Urban + log_Altitude + ULTISOLES + ULTISOLES_INCEPTISOLES  
                  + ENTISOLES + INCEPTISOLES + INCEPTISOLES_ANDISOLES, data = umf_3, K = 50, engine = "R")
large_23 <- occuRN(~time.of.day.factor ~Urban + log_Altitude + FLII, data = umf_3, K = 50, engine = "R")
large_24 <- occuRN(~time.of.day.factor ~Urban + FLII + ULTISOLES + ULTISOLES_INCEPTISOLES  
                  + ENTISOLES + INCEPTISOLES + INCEPTISOLES_ANDISOLES, data = umf_3, K = 50, engine = "R")

large_25 <- occuRN(~time.of.day.factor ~FLII + log_Altitude + ULTISOLES + ULTISOLES_INCEPTISOLES  
                  + ENTISOLES + INCEPTISOLES + INCEPTISOLES_ANDISOLES, data = umf_3, K = 50, engine = "R")

# 4 covariates
large_26 <- occuRN(~time.of.day.factor ~Trees + FLII + log_Altitude + ULTISOLES + ULTISOLES_INCEPTISOLES  
                  + ENTISOLES + INCEPTISOLES + INCEPTISOLES_ANDISOLES, data = umf_3, K = 50, engine = "R")
large_27 <- occuRN(~time.of.day.factor ~Trees + FLII + Urban + ULTISOLES + ULTISOLES_INCEPTISOLES  
                  + ENTISOLES + INCEPTISOLES + INCEPTISOLES_ANDISOLES, data = umf_3, K = 50, engine = "R")
large_28 <- occuRN(~time.of.day.factor ~Trees + FLII + log_Altitude + Urban, data = umf_3, K = 50, engine = "R")
large_29 <- occuRN(~time.of.day.factor ~Trees + Urban + log_Altitude + ULTISOLES + ULTISOLES_INCEPTISOLES  
                  + ENTISOLES + INCEPTISOLES + INCEPTISOLES_ANDISOLES, data = umf_3, K = 50, engine = "R")

large_30 <- occuRN(~time.of.day.factor ~Urban + FLII + log_Altitude + ULTISOLES + ULTISOLES_INCEPTISOLES  
                  + ENTISOLES + INCEPTISOLES + INCEPTISOLES_ANDISOLES, data = umf_3, K = 50, engine = "R")
large_31 <- occuRN(~time.of.day.factor ~Urban + FLII + Trees + ULTISOLES + ULTISOLES_INCEPTISOLES  
                  + ENTISOLES + INCEPTISOLES + INCEPTISOLES_ANDISOLES, data = umf_3, K = 50, engine = "R")

# 5 covariates
large_32 <- occuRN(~time.of.day.factor ~ Trees + Urban + log_Altitude + FLII + ULTISOLES + 
    ULTISOLES_INCEPTISOLES + ENTISOLES + INCEPTISOLES + INCEPTISOLES_ANDISOLES, data = umf_3, K = 50, engine = "R")
```

```
# create model table
table <- list(large_1, large_2, large_3, large_4, large_5, large_6, large_7, large_8, large_9, large_10,
     large_11, large_12, large_13, large_14, large_15, large_16, large_17, large_18, large_19, 
     large_20, large_21, large_22, large_23, large_24, large_25, large_26, large_27, large_28, large_29,
     large_30, large_31, large_32) %>% 
  map(~.@AIC) %>% 
  set_names(nm = c(" ~Trees", 
     " ~Urban", 
     " ~log(Altitude)", 
     " ~FLII",
     " ~soil order", 
     " ~Trees + Urban", 
     " ~Trees + log(Altitude)", 
     " ~Trees + FLII", 
     " ~Trees + soil order",
     " ~Urban + FLII", 
     " ~Urban + log(Altitude)", 
     " ~Urban + soil order", 
     " ~log(Altitude) + FLII", 
     " ~log(Altitude) + soil order", 
     " ~FLII + soil order", 
     " ~Trees + Urban + log(Altitude)", 
     " ~Trees + Urban + FLII", 
     " ~Trees + Urban + soil order", 
     " ~Trees + FLII + log(Altitude)",
     " ~Trees + FLII + soil order", 
     " ~Trees + log(Altitude) + soil order",
     " ~Urban + log(Altitude) + soil order",
     " ~Urban + log(Altitude) + FLII",
     " ~Urban + FLII + soil order",
     " ~FLII + log(Altitude) + soil order", 
     " ~Trees + FLII + log(Altitude) + soil order",
     " ~Trees + FLII + Urban + soil order",
     " ~Trees + FLII + log(Altitude) + Urban",
     " ~Trees + Urban + log(Altitude) + soil order",
     " ~Urban + FLII + log(Altitude) + soil order",
     " ~Urban + FLII + Trees + soil order",
     " ~Trees + Urban + log(Altitude) + FLII + soil order")) %>% 
  bind_rows() %>% 
  pivot_longer(cols = 1:32, names_to = "Model", values_to = "AIC") %>% 
  arrange(AIC) %>% 
  mutate(delta = round(AIC -(AIC[1]),2),
         delta = replace_na(delta, 0.00)) 

# create full model flextable to export to word
my_ft <- flextable(table, 
  col_keys = c("Model", "AIC", "delta")) %>% 
  width(width = c(4,1,1)) %>% 
  padding(padding = 1) %>% 
  theme_vanilla()

# save
save_as_docx(my_ft, path = "./model_table.docx")
```

##### Goodness of fit analysis

Taking the top model of 1km with all covariates we will now look at
the goodness-of-fit using the parboot function from
`unmarked`

```
# Run the parboot function with the default statistic: sum of squared residuals
pb <- parboot(large_26, nsim=1000, report=1, seed = set.seed(345), ncores = 6)
```

```
## t0 = 466.3791
```

```
# plot to see the results
pb_df <- pb@t.star %>% 
  as_tibble() %>%
  rename(SSE = 1) 

pb_sum <- pb_df %>% 
  summarise(mean = mean(SSE),
            SD = sd(SSE),
            mean_diff = mean(pb@t0 - SSE),
            SD_diff = sd(pb@t0 - SSE)) %>% 
  mutate(t0 = as.numeric(pb@t0))

label_1 <- paste0("Model SSE: ", round(pb_sum$t0,2))
label_2 <- paste0("Mean Bootstrapped SSE: ", round(pb_sum$mean, 2))

ggplot() +
  geom_histogram(data = pb_df, aes(x = SSE)) +
  geom_vline(aes(xintercept = 466)) +
  geom_vline(data = pb_sum, aes(xintercept = mean), linetype = "dashed", colour = "red") +
  scale_x_continuous(limits = c(370, 600), expand = c(0,0)) +
  scale_y_continuous(expand = c(0.02,0)) +
  theme(panel.grid.major.x = element_blank(),
        panel.grid.minor.y = element_blank(),
        panel.grid.major.y = element_line(colour = "gray90"),
        panel.background = element_blank(),
        panel.border = element_rect(fill = NA, colour = "gray40"),
        axis.ticks = element_line(colour = "gray40"),
        axis.title = element_text(size = 14, colour = "gray40"),
        axis.text = element_text(size = 12, colour = "gray40"),
        plot.margin = margin(5,5,5,5, unit = "pt")) +
  geom_textbox(aes(x = 550, y = 88, label = label_2, halign = 1), 
               width = unit(6,"cm"), fill = "mistyrose", colour = "red", 
               text.colour = "black", size = 4) +
  geom_textbox(aes(x = 550, y = 97, label = label_1, halign = 1),  
               width = unit(6,"cm"), size = 4) +
  ylab("Count") +
  xlab("SSE")
```

```
ggsave(filename = "./model_GOF_his.png", width = 35, height = 20, dpi=300, units = "cm")
```

##### Estimated population at sites

```
# calculate site totals
site_preds <- predict(large_26,
                    type = "state",
                    newdata = umf_3,
                    na.rm = TRUE) %>% 
  bind_cols(cov_time_factor$site, .) %>% 
  rename(site = 1) %>% 
  mutate(Type = "Predict") %>% 
  left_join(., read_csv("./Data/site coordinates UTM.csv"), by = "site")

site_preds_ranef <- ranef(large_32, 1000) %>% 
        bup(., stat = "mean") %>% 
        as_tibble() %>% 
        rename(Predicted = value) %>% 
        mutate(Predicted = round(Predicted, 3)) %>% 
  bind_cols(., ranef(large_32, 1000) %>%  
        confint(., level=0.9) %>% 
          as_tibble() %>% 
          rename(lower = "5%",
                 upper = "95%"))  %>% 
  bind_cols(cov_time_factor$site, .) %>%
  rename(site = 1) %>%
  left_join(., read_csv("./Data/site coordinates UTM.csv"), by = "site") %>% 
  mutate(Type = "EBE") %>% 
  left_join(., read_csv("./Data/covariate_site_grid_extended_10000m.csv") %>% rename(site = .id), by = "site")

site_preds_all <- site_preds_ranef %>% 
  bind_rows(., site_preds %>% dplyr::select(-SE))
```

###### Standard Errors

Calculate the standard errors on the total population estimate across
the sites.

```
# first get the covariate coefficients:
state_estimates <- as.vector(large_26@estimates@estimates[["state"]]@estimates)

# and the covariance martrix
state_matrix <- large_26@estimates@estimates[["state"]]@covMat

# use deltavar method
Var_hat<-deltavar(fun = sum(exp(B0 + B1 * site_covs[[2]]$Trees + 
                                  B2 * site_covs[[2]]$FLII + 
                                  B3 * site_covs[[2]]$log_Altitude + 
                                  B4 * site_covs[[2]]$ULTISOLES +
                                  B5 * site_covs[[2]]$ULTISOLES_INCEPTISOLES +
                                  B6 * site_covs[[2]]$ENTISOLES +
                                  B7 * site_covs[[2]]$INCEPTISOLES + 
                                  B8 * site_covs[[2]]$INCEPTISOLES_ANDISOLES)), 
                  meanval = c(B0 = state_estimates[1], B1 = state_estimates[2], 
                              B2 = state_estimates[3], B3 = state_estimates[4],
                              B4 = state_estimates[5], B5 = state_estimates[6], 
                              B6 = state_estimates[7], B7 = state_estimates[8],
                              B8 = state_estimates[9]),
                  Sigma = state_matrix, verbose = T)
```

```
## value of derivs:
##         [,1]     [,2]     [,3]      [,4]     [,5]      [,6]      [,7]      [,8]
## [1,] 208.476 158.8726 56.05494 -95.99253 168.3394 -9.403452 -30.76168 -69.60616
##           [,9]
## [1,] -56.74505
```

```
# calculate the SE
SE<-sqrt(Var_hat)

cat("\nTotal population:",sum(site_preds$Predicted), "+/-", SE, "SE\n")
```

```
## 
## Total population: 208.476 +/- 30.36298 SE
```

###### Plot site level abundance

Have a look at the number of individuals estimated at each site in a
map

```
# get a squared grid for the sites
pc_initial <- read_csv("./Data/PC location 2020.csv") %>% 
  filter(Site > 0) %>% 
  rename(id = Site)

# add this to the data from sites
site_map <- site_preds_all %>%  
  left_join(., pc_initial, by = "id") %>% 
  st_as_sf(coords = c("x","y"), crs = 32616)

# load the studysite
studysite <- st_read("./Data/grid_ss.shp") %>% 
  st_set_crs(4326) %>% 
  st_transform(32616)
```

```
## Reading layer `grid_ss' from data source 
##   `H:\PhD\1. Chapters\R\TTD\Data\grid_ss.shp' using driver `ESRI Shapefile'
## Simple feature collection with 1 feature and 2 fields
## Geometry type: POLYGON
## Dimension:     XY
## Bounding box:  xmin: -84.4 ymin: 10.1 xmax: -83.17764 ymax: 10.99399
## Geodetic CRS:  GCS_unknown
```

```
# load flii data to use as nice base
flii_data <- raster("./Data/range_flii_rescale_crop.tif") 
flii_data <- mask(crop(flii_data, studysite), studysite)
  
flii_df <- as(flii_data, "SpatialPixelsDataFrame")
flii_df_plot <- as.data.frame(flii_df)
colnames(flii_df_plot) <- c("FLII", "x", "y") 

flii_df_plot <- flii_df_plot %>% 
  mutate(ints = floor(FLII),
         category = ifelse(ints <6, "Low", "Medium"))

# plot with radius equal to estimated pop per siite
ggplot() +
  geom_raster(data = flii_df_plot, aes(x = x, y = y, fill = category), alpha = 0.5) +
  scale_fill_manual(values = c("khaki","darkorange4")) +
  geom_sf(data = site_map %>%  filter(Type == "EBE"), mapping = aes(size = Predicted), fill = "darkgreen",
             colour = "darkgreen", shape = 21) +
  geom_sf_text(data = site_map, aes(label = id), 
               colour = "gray40", size = 4, hjust = 0, nudge_x = 2500) +
  annotation_north_arrow(location = "tr", which_north = "true", 
        pad_x = unit(0.2, "cm"), pad_y = unit(0.2, "cm"),
        width = unit(1, "cm"), height = unit(1, "cm"),
        style = north_arrow_orienteering) +
    annotation_scale(bar_cols = c("gray40", "white"), height = unit(0.2, "cm")) +
  scale_alpha_continuous(name="Estimated\nabundance") +
  scale_size_continuous(name="Estimated\nabundance") +
  scale_x_continuous(expand = c(0,0)) +
  scale_y_continuous(expand = c(0,0)) +
  theme(panel.grid.major.x = element_blank(),
        panel.grid.minor.y = element_blank(),
        panel.grid.major.y = element_line(colour = "gray80"),
        panel.background = element_blank(),
        panel.border = element_rect(fill = NA, colour = "gray40"),
        axis.ticks = element_line(colour = "gray40"),
        axis.title = element_text(size = 12, colour = "gray40"),
        axis.text = element_text(size = 10),
        legend.title = element_text(size = 10, colour = "gray40"),
        legend.text = element_text(size = 10, colour = "gray40"),
        legend.key = element_blank(),
        plot.margin = margin(5,5,5,5, unit = "pt"),
        legend.position = "right") +
  guides(fill = guide_legend(title = "FLII\ncategory")) +
  xlab("Long") +
  ylab("Lat")
```

```
ggsave(filename = "./Site level abundance FLII Low-Med.png", width = 25, height = 15, dpi=300, units = "cm")
```

###### Coeefficent plots

Now I am going to create coefficient plots for the top model

```
model_coef <- as_tibble(t(coef(large_26))) %>% 
  pivot_longer(cols = 1:ncol(.), names_to = "Variable", values_to = "Value") %>% 
  mutate(model = "large_26",
         covariate = str_remove(str_extract(string = Variable,
                                               pattern = "(?<=\\().*(?=\\))"), "ORDENES"),
         covariate = str_remove(covariate, "time.of.day.factor"),
         covariate = ifelse(covariate == "log_Altitude", "log(Altitude)", covariate),
         state = sub("\\(.*","", Variable),
         state = ifelse(state == "lam", "Abundance model", "Detection model")) 

# get SE and confidence intervals of the transformed coefficients
model_confint_coef <- as_tibble(t(confint(large_26, type = "state"))) %>% 
  bind_cols(., as_tibble(t(confint(large_26, type = "det")))) %>% 
  bind_cols(., c("lower","upper")) %>% 
  rename(Interval = ncol(.)) %>% 
  pivot_longer(cols = 1:(ncol(.)-1), names_to = "Variable", values_to = "Value") %>% 
  pivot_wider(names_from = "Interval", values_from = "Value") %>% 
  left_join(model_coef, ., by = "Variable") %>% 
  bind_cols(., sqrt(diag(vcov(large_26))) %>% 
              as_tibble() %>% 
              rename(SE = value)) %>% 
  mutate(order = ifelse(covariate == "Int" & state  == "Abundance mode", 1, 
                        ifelse(covariate %in% c("Urban","Trees","FLII"), 4,
                        ifelse(grepl("SOLES", covariate), 2, 
                              ifelse(grepl("time.of.day", Variable), 6,
                                       ifelse(covariate == "log(Altitude)", 3, 
                                              ifelse(covariate == "Int" & state  == "Detection model", 5, 4)))))),
         names = ifelse(order == 1, "Intercept", 
                        ifelse(order == 2, "Soil order", 
                               ifelse(order == 3, "log(Altitude)", 
                                      ifelse(order == 4, "Habitat", "Time"))))) %>% 
  arrange(order) %>% 
  mutate(val = row_number(),
         covariate = ifelse(order == 2, gsub("(\\b[A-Z])[^A-Z]+", "\\1", 
                                             str_to_title(gsub("_", " ", covariate, fixed=TRUE)), perl = TRUE),
                            covariate))

ggplot(model_confint_coef) +
  geom_point(aes(x = Value, y = reorder(covariate, order), colour = names), size = 4) +
  geom_linerange(aes(xmin = lower, xmax = upper, y = covariate, colour = names)) +
  geom_vline(aes(xintercept = 0), colour = "black", linetype = "dotted") +
  facet_wrap(~state, scales = "free") +
  theme(panel.grid.major.x = element_blank(),
        panel.grid.minor.y = element_blank(),
        panel.grid.major.y = element_line(colour = "gray80"),
        panel.background = element_blank(),
        panel.border = element_rect(fill = NA, colour = "gray40"),
        axis.ticks = element_line(colour = "gray40"),
        axis.title.y = element_text(size = 12, colour = "gray40"),
        axis.text = element_text(size = 10),
        axis.title.x = element_blank(),
        legend.title = element_text(size = 12, colour = "gray40"),
        legend.text = element_text(size = 10, colour = "gray40"),
        legend.key = element_blank(),
        plot.margin = margin(0,0,0,0, unit = "pt"),
        strip.background =element_rect(colour = "gray40", fill = "honeydew2"),
        strip.text = element_text(size = 12),
        legend.position = "right",
        strip.text.y.right = element_text(angle = 0)) +
  guides(colour = guide_legend(title = "Covariate Type")) +
  ylab("Covariate")
```

```
ggsave(filename = "./RN Model coefficient plot.png", width = 25, height = 15, dpi=300, units = "cm")
```

We now need to get the scale values for each covariate so we can
apply this to the covariates when we got to predict over the whole
range

```
model_scale <- c(attr(site_covs[[2]]$Trees, 'scaled:scale'),
                 attr(site_covs[[2]]$FLII, 'scaled:scale'), 
                 attr(site_covs[[2]]$log_Altitude, 'scaled:scale'),
                 attr(site_covs[[2]]$ULTISOLES, 'scaled:scale'),
                 attr(site_covs[[2]]$ULTISOLES_INCEPTISOLES, 'scaled:scale'),
                 attr(site_covs[[2]]$INCEPTISOLES, 'scaled:scale'),
                 attr(site_covs[[2]]$INCEPTISOLES_ANDISOLES, 'scaled:scale'),
                 attr(site_covs[[2]]$ENTISOLES, 'scaled:scale'))

model_center <- c(attr(site_covs[[2]]$Trees, 'scaled:center'), 
                 attr(site_covs[[2]]$FLII, 'scaled:center'), 
                 attr(site_covs[[2]]$log_Altitude, 'scaled:center'),
                 attr(site_covs[[2]]$ULTISOLES, 'scaled:center'),
                 attr(site_covs[[2]]$ULTISOLES_INCEPTISOLES, 'scaled:center'),
                 attr(site_covs[[2]]$INCEPTISOLES, 'scaled:center'),
                 attr(site_covs[[2]]$INCEPTISOLES_ANDISOLES, 'scaled:center'),
                 attr(site_covs[[2]]$ENTISOLES, 'scaled:center'))

names <- c("Trees", "FLII","log_Altitude", "ULTISOLES", "ULTISOLES_INCEPTISOLES", 
           "INCEPTISOLES","INCEPTISOLES_ANDISOLES", "ENTISOLES")

model_transform <- bind_cols(as_tibble(names), as_tibble(model_scale), as_tibble(model_center)) %>% 
  set_names(nm = c("covariate","sd", "mean"))
```

##### Predict over the whole range

```
# Read in data for the whole of the GGMs historic range
ggPred <- read_csv("./Data/covariate_extended_full_grid_10km.csv") %>% 
  mutate(log_Altitude = log10(Altitude)) %>% 
  dplyr::select(.id, Trees, Urban, log_Altitude, FLII, ULTISOLES, ULTISOLES_INCEPTISOLES, 
           INCEPTISOLES, INCEPTISOLES_ANDISOLES, ENTISOLES) %>% 
  pivot_longer(cols = 2:ncol(.), names_to = "covariate", values_to = "values")%>% 
  left_join(., model_transform, by = "covariate") %>% 
  mutate(final = (values - mean)/sd) %>%
  dplyr::select(.id, covariate, final) %>% 
  pivot_wider(names_from = "covariate", values_from = "final") %>% 
  data.frame(.)

# find grid squares that are within the ebird range
pred_actual_id_mr <- st_read("./Data/range_grid_10km.shp") %>% rename(.id = X_id) %>% 
  st_intersection(., st_read("./Data/ebird_ggm_range_mr.shp"))
```

```
## Reading layer `range_grid_10km' from data source 
##   `H:\PhD\1. Chapters\R\TTD\Data\range_grid_10km.shp' using driver `ESRI Shapefile'
## Simple feature collection with 200 features and 1 field
## Geometry type: POLYGON
## Dimension:     XY
## Bounding box:  xmin: 663374.8 ymin: 1028424 xmax: 983374.8 ymax: 1228424
## Projected CRS: WGS 84 / UTM zone 16N
## Reading layer `ebird_ggm_range_mr' from data source 
##   `H:\PhD\1. Chapters\R\TTD\Data\ebird_ggm_range_mr.shp' using driver `ESRI Shapefile'
## Simple feature collection with 1 feature and 9 fields
## Geometry type: MULTIPOLYGON
## Dimension:     XY
## Bounding box:  xmin: 772469.9 ymin: 1063023 xmax: 981028.5 ymax: 1217290
## Projected CRS: WGS 84 / UTM zone 16N
```

```
pred_actual_id_raw <- st_read("./Data/range_grid_10km.shp") %>% rename(.id = X_id) %>% 
  st_intersection(., st_read("./Data/ebird_ggm_range_raw.shp"))
```

```
## Reading layer `range_grid_10km' from data source 
##   `H:\PhD\1. Chapters\R\TTD\Data\range_grid_10km.shp' using driver `ESRI Shapefile'
## Simple feature collection with 200 features and 1 field
## Geometry type: POLYGON
## Dimension:     XY
## Bounding box:  xmin: 663374.8 ymin: 1028424 xmax: 983374.8 ymax: 1228424
## Projected CRS: WGS 84 / UTM zone 16N
## Reading layer `ebird_ggm_range_raw' from data source 
##   `H:\PhD\1. Chapters\R\TTD\Data\ebird_ggm_range_raw.shp' using driver `ESRI Shapefile'
## Simple feature collection with 1 feature and 9 fields
## Geometry type: MULTIPOLYGON
## Dimension:     XY
## Bounding box:  xmin: 738168.5 ymin: 1053247 xmax: 984078.3 ymax: 1218642
## Projected CRS: WGS 84 / UTM zone 16N
```

```
# ebird isolated sightings
sighting <- st_read("./Data/range_grid_10km.shp") %>% rename(.id = X_id) %>% 
  st_intersection(., st_read("./Data/ebird_isolated_sighting.shp"))
```

```
## Reading layer `range_grid_10km' from data source 
##   `H:\PhD\1. Chapters\R\TTD\Data\range_grid_10km.shp' using driver `ESRI Shapefile'
## Simple feature collection with 200 features and 1 field
## Geometry type: POLYGON
## Dimension:     XY
## Bounding box:  xmin: 663374.8 ymin: 1028424 xmax: 983374.8 ymax: 1228424
## Projected CRS: WGS 84 / UTM zone 16N
## Reading layer `ebird_isolated_sighting' from data source 
##   `H:\PhD\1. Chapters\R\TTD\Data\ebird_isolated_sighting.shp' 
##   using driver `ESRI Shapefile'
## Simple feature collection with 5 features and 10 fields
## Geometry type: POINT
## Dimension:     XY
## Bounding box:  xmin: 743897.8 ymin: 1057716 xmax: 945066.7 ymax: 1206249
## Projected CRS: WGS 84 / UTM zone 16N
```

```
# filter to size 
ggEbird_mr <- ggPred %>% 
  filter(.id %in% pred_actual_id_mr$.id)

# filter to size
ggEbird_raw <- ggPred %>% 
  filter(.id %in% pred_actual_id_raw$.id)

# filter to size
ggIsolated <- ggPred %>% 
  filter(.id %in% sighting$.id)

# turn into a list
pred_data <- list(ggPred, ggEbird_raw, ggEbird_mr, ggIsolated)

# now predict:
occuPred <- pred_data %>%  
  bind_rows() %>% 
  predict(large_26, type = "state", newdata = ., na.rm = TRUE, inf.rm = TRUE)

# join predictions to the corresponding grid ids for each abundance area
pred_final <- pred_data %>% 
  map2(., c("Full", "ebird_mr", "ebird_raw","isolated"), ~.x %>% mutate(Type = .y)) %>% 
  bind_rows() %>%
  dplyr::select(Type, .id) %>% 
  bind_cols(., occuPred) %>% 
  left_join(sf::st_read("./Data/range_grid_10km.shp") %>% rename(.id = X_id), ., by = ".id")
```

```
## Reading layer `range_grid_10km' from data source 
##   `H:\PhD\1. Chapters\R\TTD\Data\range_grid_10km.shp' using driver `ESRI Shapefile'
## Simple feature collection with 200 features and 1 field
## Geometry type: POLYGON
## Dimension:     XY
## Bounding box:  xmin: 663374.8 ymin: 1028424 xmax: 983374.8 ymax: 1228424
## Projected CRS: WGS 84 / UTM zone 16N
```

```
pred_sum <- pred_final %>% 
  st_drop_geometry() %>% 
  group_by(Type) %>% 
  summarise(total = sum(Predicted)) %>% 
  mutate(SE = 1) %>% # this has to be numeric so the loop can change to the SE
  arrange(desc(total))

# use deltavar method again. We are using the same coefficients as before but changing the covariate values:

for (i in seq_along(pred_data)) {
Var_hat<-deltavar(fun = sum(exp(B0 + B1 * pred_data[[i]]$Trees + 
                                  B2 * pred_data[[i]]$FLII + 
                                  B3 * pred_data[[i]]$log_Altitude + 
                                  B4 * pred_data[[i]]$ULTISOLES +
                                  B5 * pred_data[[i]]$ULTISOLES_INCEPTISOLES +
                                  B6 * pred_data[[i]]$INCEPTISOLES + 
                                  B7 * pred_data[[i]]$INCEPTISOLES_ANDISOLES +
                                  B8 * pred_data[[i]]$ENTISOLES)), 
                  meanval = c(B0 = state_estimates[1], B1 = state_estimates[2], 
                              B2 = state_estimates[3], B3 = state_estimates[4],
                              B4 = state_estimates[5], B5 = state_estimates[6], 
                              B6 = state_estimates[7], B7 = state_estimates[8],
                              B8 = state_estimates[9]),
                  Sigma = state_matrix, verbose = T)

  # calculate the SE
  pred_sum[i,3] <- sqrt(Var_hat)
}
```

```
## value of derivs:
##          [,1]     [,2]      [,3]      [,4]     [,5]      [,6]      [,7]
## [1,] 762.9706 374.3044 -14.83387 -184.8508 1046.286 -495.1678 -98.97249
##           [,8]      [,9]
## [1,] -275.4994 -152.4894
## value of derivs:
##         [,1]     [,2]     [,3]      [,4]     [,5]     [,6]      [,7]      [,8]
## [1,] 517.779 362.3905 123.8973 -186.7958 698.3841 -287.288 -103.3599 -178.5273
##           [,9]
## [1,] -113.1544
## value of derivs:
##          [,1]     [,2]     [,3]      [,4]     [,5]      [,6]      [,7]     [,8]
## [1,] 402.1687 291.2548 115.9552 -138.3584 534.3394 -185.9988 -76.55763 -132.804
##           [,9]
## [1,] -101.1361
## value of derivs:
##          [,1]     [,2]      [,3]      [,4]     [,5]      [,6]     [,7]     [,8]
## [1,] 52.42465 30.58612 -5.178162 -43.87361 55.92752 -45.93051 2.306786 -20.7337
##           [,9]
## [1,] -7.129886
```

```
cat("Predicted population size within smoothed ebird range = ", pred_sum$total[2], "+/-", pred_sum$SE[2], "\n")
```

```
## Predicted population size within smoothed ebird range =  637.9217 +/- 79.81795
```

```
cat("Predicted population size in isolated ebird areas = ", pred_sum$total[4], "+/-", pred_sum$SE[4], "\n")
```

```
## Predicted population size in isolated ebird areas =  49.34231 +/- 11.18363
```

```
cat("Predicted population size within raw ebird range = ", pred_sum$total[3], "+/-", pred_sum$SE[3], "\n")
```

```
## Predicted population size within raw ebird range =  485.6576 +/- 60.90663
```

```
cat("Potential population size in CR = ", pred_sum$total[1], "+/-", pred_sum$SE[1], "\n")
```

```
## Potential population size in CR =  883.0875 +/- 128.1633
```

###### Plot whole range estimate

```
# get just outline of ebird range
ebird_raw_outline <- pred_final %>% 
  filter(Type == "ebird_raw") %>% 
  st_union(.)

ebird_mr_outline <- pred_final %>% 
  filter(Type == "ebird_mr") %>% 
  st_union(.)

# get the range outline 
range_outline <- pred_final %>% 
  filter(Type == "Full") %>% 
  st_union(.)

# load protected areas and clip to the historical range:
PA <- st_read("./Data/Areassilvestresprotegidas2014crtm05.shp") %>% 
  st_transform(crs= 32616) %>% 
  st_intersection(., range_outline) %>% 
  mutate(Type = ifelse(grepl("MIXTO", NOMBRE_), "Mixed", "Full"))
```

```
## Reading layer `Areassilvestresprotegidas2014crtm05' from data source 
##   `H:\PhD\1. Chapters\R\TTD\Data\Areassilvestresprotegidas2014crtm05.shp' 
##   using driver `ESRI Shapefile'
## Simple feature collection with 505 features and 6 fields
## Geometry type: MULTIPOLYGON
## Dimension:     XY
## Bounding box:  xmin: 207669 ymin: 906446.4 xmax: 658789.3 ymax: 1241133
## Projected CRS: Proyección CRTM05
```

```
# load the biological corridors and clip to the historical range
BC <- st_read("./Data/CorredoresBiologicos2009_SINAC.shp") %>% 
  st_transform(crs= 32616) %>% 
  filter(NOMBRE___ == "SAN JUAN LA SELVA") %>% 
  st_intersection(., range_outline) %>% 
  mutate("Biological\nCorridor" = "SJLA")
```

```
## Reading layer `CorredoresBiologicos2009_SINAC' from data source 
##   `H:\PhD\1. Chapters\R\TTD\Data\CorredoresBiologicos2009_SINAC.shp' 
##   using driver `ESRI Shapefile'
## Simple feature collection with 192 features and 3 fields
## Geometry type: MULTIPOLYGON
## Dimension:     XY
## Bounding box:  xmin: 295652.5 ymin: 925977.3 xmax: 658794 ymax: 1223319
## Projected CRS: Proyección CRTM05
```

```
# plot
ggplot() +
  geom_sf(pred_final %>% filter(Type == "Full"), mapping = aes(fill = Predicted), colour = NA) + 
  geom_sf(range_outline, mapping = aes(), colour = "gray40", fill = NA) +
  scale_fill_gradient(low = "beige", high = "darkgreen") +
  new_scale("fill") +
  geom_sf(BC, mapping = aes(fill = "Biological\nCorridor"), colour = "gray40", alpha = 0.1) +
  geom_sf_pattern(PA, mapping = aes(pattern = Type, pattern_colour = Type, pattern_size = Type),  
                  colour = "gray50", fill = NA, alpha = 0.5,
                  pattern_spacing = 0.015, 
                  pattern_alpha = 0.5) +
  scale_pattern_size_manual(values = c(0.8, 1.5)) +
  scale_pattern_colour_manual(values = c("gray50", "gray20")) +
  geom_sf(ebird_mr_outline, mapping = aes(colour = "ebird range"), 
          fill = NA, size = 1, show.legend = "polygon")+
  geom_sf(sighting, mapping = aes(), colour = "red") +
  guides(colour = guide_legend(title = "", nrow = 1), fill = guide_legend(title = ""), 
         pattern = guide_legend(title = "PA Type"), pattern_size = guide_legend(title = "PA Type"),
         pattern_colour = guide_legend(title = "PA Type")) +
  annotation_north_arrow(location = "tr", which_north = "true", 
        pad_x = unit(0.2, "cm"), pad_y = unit(0.2, "cm"),
        width = unit(1, "cm"), height = unit(1, "cm"),
        style = north_arrow_orienteering) +
  annotation_scale(bar_cols = c("gray40", "white"), height = unit(0.2, "cm")) +
  scale_fill_manual(values = "red") +
  theme(panel.grid.major.x = element_blank(),
        panel.grid.minor.y = element_blank(),
        panel.grid.major.y = element_line(colour = "gray80"),
        panel.background = element_blank(),
        panel.border = element_rect(fill = NA, colour = "gray40"),
        axis.ticks = element_line(colour = "gray40"),
        axis.title = element_text(size = 12, colour = "gray40"),
        axis.text = element_text(size = 10),
        legend.title = element_text(size = 10, colour = "gray40"),
        legend.text = element_text(size = 10, colour = "gray40"),
        legend.key = element_blank(),
        plot.margin = margin(5,5,5,5, unit = "pt"),
        legend.position = "right") +
  xlab("Long") +
  ylab("Lat")
```

```
ggsave(filename = "./RN Predicted Abundance PA ebird smoothed Map points.png", width = 25, height = 15, dpi=300, units = "cm")
```
